# Transcriptional regulation of Dyskerin *via* canonical WNT signaling modulates sphingolipid biosynthesis and drives colorectal cancer

**DOI:** 10.1101/2023.07.18.549480

**Authors:** Shivansh Nigam, Umar K. Khan, Ayush Praveen, Akshay Shendre, Shannon Carskadon, Abhimanyu Kapoor, Anjali Tiwari, Abhijit Chandra, Nallasivam Palanisamy, Bushra Ateeq

**Affiliations:** Molecular Oncology Laboratory, Department of Biological Sciences and Bioengineering, Indian Institute of Technology Kanpur, Kanpur, UP, 208016, INDIA; Department of Urology, Vattikutti Urology Institute, Henry Ford Health System, Detroit, MI, 48202, USA; Regency Hospital, Kanpur, UP, 208005, INDIA; Department of Surgical Gastroenterology, King George Medical University, Lucknow, UP, 226003, INDIA; The Mehta Family Center for Engineering in Medicine, Indian Institute of Technology Kanpur, Kanpur, UP, 208016, INDIA; Centre of Excellence for Cancer, Gangwal School of Medical Sciences and Technology, Indian Institute of Technology Kanpur, Kanpur, 208016, UP, INDIA

## Abstract

Targeting EGFR has been effective in RAS/RAF wild-type colorectal cancer (CRC) patients. However, residual tumor relapses, necessitating the importance of biomarker-guided novel therapeutics. We show elevated DKC1 in ∼88% of CRC patients with poor recurrence-free survival. Clinically, *DKC1-*positive patients exhibit similarity with CMS2 class, the canonical subtype with active WNT signaling. We show functional significance of DKC1 in cell proliferation, stemness, DNA repair, and survival. Further, mice bearing *DKC1* knockdown xenografts show ∼81% reduction in tumor burden. Mechanistically, WNT/β-catenin signaling orchestrates *DKC1* expression, then, DKC1/SOX2 complex regulates *SGPP2*, modulating sphingolipids metabolism. Downregulation of DKC1 in CRC lead to reduced SGPP2 levels leading to dysregulation of sphingolipid biosynthesis. Of note, *DKC1*-high CRC patients show accumulation of ceramides, namely C23 and C24, signifying their utility in diagnosis. Collectively, we delineate the mechanistic circuitry involved in DKC1-mediated CRC progression, propose ceramides as biomarker, and underscore WNT-based therapeutics for DKC1-positive patients.

## INTRODUCTION

The epidermal growth factor receptor (EGFR) belonging to the receptor tyrosine kinase super- family is highly expressed in metastatic CRC and has been used as a therapeutic target^1^. Besides EGFR, aberrantly active WNT signaling play a causal role in the majority of colorectal cancer (CRC) patients, however targeting the WNT pathway poses significant challenges^2–4^. WNT inhibitors targeting the catalytic production of WNT ligands, antagonists for WNT receptors, or inhibitors of transcriptional messengers such as β-catenin have been rigorously exploited against aggressive cancers, some of these are in clinical trials^5^. Moreover, a significant fraction of CRC patients often receive multimodal treatment such as 5-fluorouracil, folinic acid, and irinotecan (FOLFIRI) or folinic acid, 5-fluorouracil, and oxaliplatin (FOLFOX), as a part of first-line therapy coupled with anti-EGFR monoclonal antibodies; however, selecting the optimum drug regimen(s) mainly relies on the molecular profiling of the tumor^6, 7^.

The molecular heterogeneity associated with CRC leads to a variable clinical course that impedes the selection of effective therapeutics, and challenges accurate evaluation of the prognostic and predictive indicators. Thus, CRC Subtyping Consortium (CRCSC) devised a criteria for categorization of patients into four consensus molecular subtypes (CMS) with distinct features: CMS1 (microsatellite instability immune, 14%) which is hypermutated, microsatellite unstable, and strong immune activation; CMS2 (canonical, 37%), epithelial, marked WNT and MYC signaling activation; CMS3 (metabolic, 13%), also epithelial and striking metabolic dysregulation; and CMS4 (mesenchymal, 23%), with prominent TGF-β activation, stromal invasion, angiogenesis, and an inflammatory, immunosuppressive phenotype^8^. Although CMS classification offers potential for clinical utility in predicting prognosis and therapy response ^9, 10^, further refinement of this classification is required for a thorough molecular understanding of these subtypes. For example, recently single-cell characterization of the CMS subtypes revealed an association of CMS2 with substantial variability and enrichment of multiple pathways regulating metabolism and cell cycle, indicating CMS2 as the highly heterogeneous subtype across various CMS subgroups^11^.

Dyskerin pseudouridine synthase 1 (*DKC1*) is mutated in an X-linked human disease known as dyskeratosis congenita^12^, and is known to be involved in pseudo-uridylation of ribosomal RNA (rRNA) and telomere maintance^13^. Moreover, high *DKC1* expression in hepatocellular and breast cancers has been associated with poor relapse-free survival and shorter breast cancer specific survival probability, respectively^14, 15^. The multipronged role of DKC1, such as in regulation of cell cycle processes and rRNA processing makes it an ideal therapeutic target for cancer^16^. Nevertheless, the multifactorial nature of DKC1 demands more insightful and elaborate elucidation of its oncogenic function, the underlying mechanism involved in its upregulation, and the associated biological pathways involved in CRC progression.

Here, we show that DKC1 levels are elevated in CRC patients, where it serves as a predictive factor. We disentangle the phenotypic and functional ramifications of DKC1-mediated downstream biological processes that operate in CRC tumorigenesis and disease progression. Our data shows the regulatory mechanism involved in its upregulation in CRC and establish its association with CMS2. Further, we show the role of DKC1 in the regulation of sphingolipid biosynthesis via SGPP2, and reveal its positive association with sphingolipids and ceramides. Notably, CRC patients with high *DKC1* tumors display an accumulation of ceramides, highlighting their utility in the diagnosis of the *DKC1*-positive sub-group. Taken together, this study provides critical insights into the mechanisms underlying DKC1–mediated CRC progression, offers ceramides as a biomarker for CRC, and provides an evidence-based therapeutic intervention to target WNT/ β-catenin signaling for the treatment of DKC1-positive CRC patients.

## RESULTS

### High DKC1 expression is associated with poor prognosis

Earlier, Firestein et al documented nine critical genes including *DKC1* by loss-of-function shRNA screens in CRC cells, wherein knockdown of these genes affected β-catenin transcriptional activity as well as cell proliferation^17^. To decipher the significance of these genes, we examined their expression using the publicly available The Cancer Genome Atlas Colorectal Adenocarcinoma (TCGA-COADREAD) dataset and found highest expression of *DKC1* in CRC tissues compared to adjacent normal tissues (**Fig. 1a**). To validate this finding, we immunostained CRC patient specimens (N=74) and high DKC1 expression was noted in ∼22% of primary tumors across all stages and grades, while medium DKC1 intensity was observed in ∼66% of the cases; only 12% showed low expression (**Fig. 1b and 1c**). Further, we also analyzed two publicly available cohorts (GSE6988 and GDC-PANCAN), and higher *DKC1* expression was noted in primary CRC compared to adjacent normal (**Fig. 1d** and Fig. S1a). Intriguingly, categorizing patients based on *KRAS* and *BRAF* mutation also exhibit high *DKC1* expression irrespective of mutational status, thus accentuating the potential of DKC1 as an important therapeutic target in a broader spectrum of patients (Fig. S1b). To examine if DKC1 holds any clinical relevance in CRC, we analyzed TCGA-COAD dataset and observed worse recurrence-free survival probability for patients with high *DKC1* expression (**Fig. 1e**). Next, we performed a comprehensive analysis using multiple publicly available CRC datasets comprising 1538 patients, which were sorted from high to low *DKC1* expression and the intergroup variability was examined using differential expression analysis between first and fourth quartiles of these patients (N=770). We found 265 genes as significantly upregulated and 51 genes exhibited low expression in *DKC1*-high vs *DKC1*-low groups (**Fig. 1f**, Supplementary table 1a and 1b), thereby a positive association of these upregulated genes with the *DKC1*-high group was noted (Fig. S1c). To identify the biological pathways associated with high *DKC1* expression, we performed gene ontology analysis and observed the enrichment of processes linked with cell cycle regulation and proliferation (**Fig. 1g**, and Supplementary table 1c and 1d).

**Figure 1:**
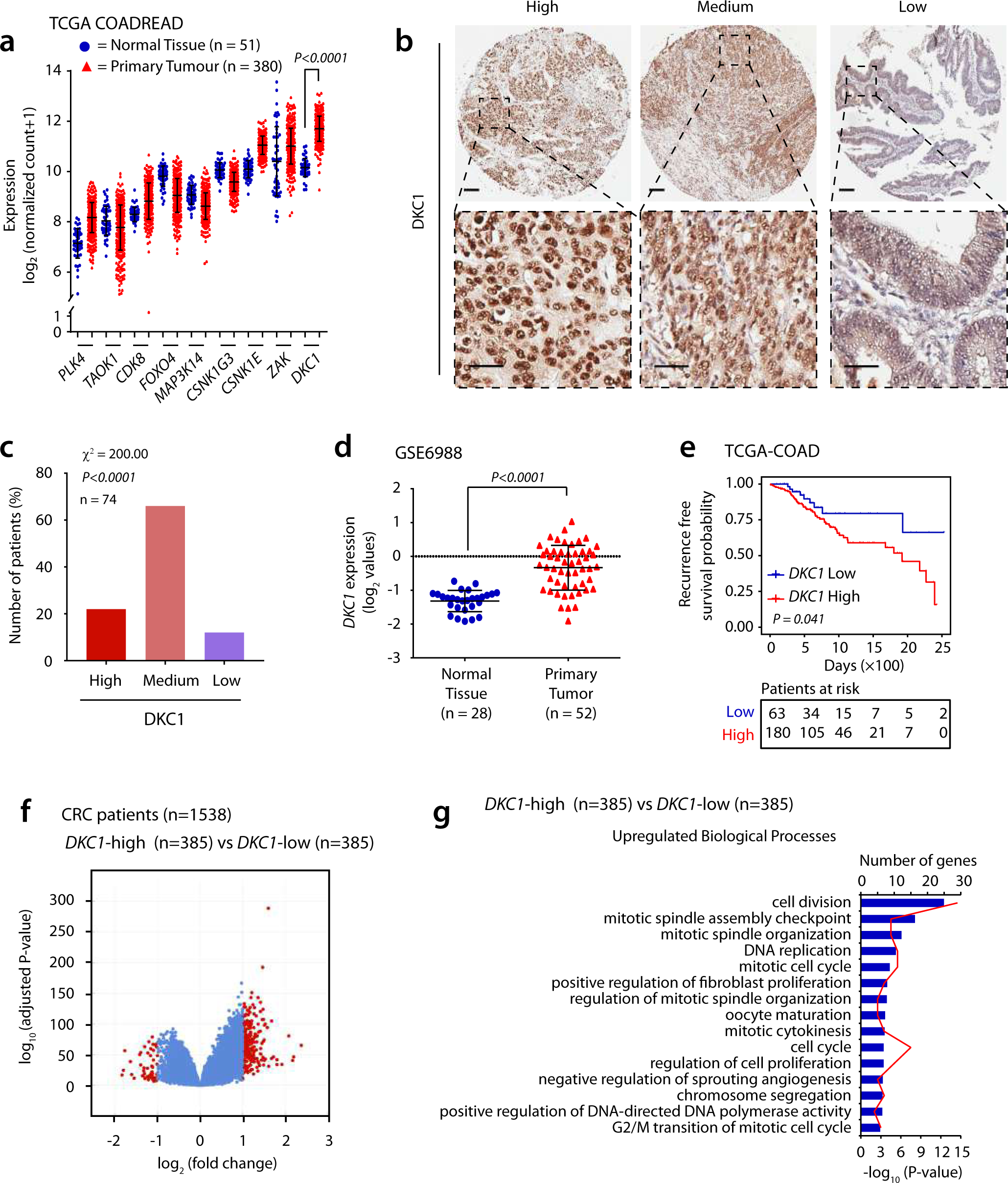
High DKC1 expression in colorectal cancer patients associate with aggressive disease and poor survival. **a,** Expression of candidate genes from shRNA screen for β-catenin activity and proliferation^17^ using publicly available data from TCGA-COADREAD cohort. **b,** Immunohistochemical (IHC) analysis of DKC1 expression using Tissue microarray comprising CRC patient specimens (N=74), scale bar for the whole section is 100 μm and for inset image is 40 μm. **c,** Quantification of IHC data for DKC1 expression for same specimens as in **b**. **d,** Expression of *DKC1* in Ki-colon cohort (GSE6988). **e,** Recurrence-free survival probability of *DKC1*-high (red) vs *DKC1*-low (blue) patients using data from TCGA COAD cohort; the number of patients at risk at a respective time point is given below. **f,** Volcano plot of differentially expressed genes using multiple CRC datasets (N=1538), data sorted as first and fourth quartile (N=770) based on *DKC1* expression; red dots represent genes with ≥ or ≤ than 2-fold change with an adjusted p-value cut-off of 0.05. **g,** Upregulated biological processes of 265 upregulated genes associated with high *DKC1* expression analyzed by DAVID functional annotation clustering ; bars (blue) represent −log10(P-value) and frequency polygon (red line) represent the number of genes. For panel **a** and **d,** Unpaired Student’s two-tailed t- test was applied. **c,** Chi-square test for comparison of frequencies was used. **e,** Cox proportional hazard analysis was performed between *DKC1*-high and *DKC1*-low groups. Data are expressed as mean ± s.e.m.

### DKC1 elicits oncogenic properties and drives CRC progression

To understand the functional significance of DKC1 in CRC, we examined the expression of DKC1-associated genes (Supplementary table 1a) in CRC cell lines, where SW620 and HT-29 (WiDr) showed higher expression of these genes (Fig. S1d). Likewise, transcriptome profile of a panel of CRC cell lines obtained from the cancer cell line encyclopedia (CCLE) revealed highest expression of *DKC1* in SW620 cells, followed by Colo320HSR and WiDr (Fig. S1e). Thus, to understand the oncogenic role of *DKC1*, the CRISPR knockout *DKC1* dependency score (Chronos and CERES, DepMap Portal) was calculated, and several CRC cell lines including SW620, Colo320HSR exhibit high dependency on DKC1 (Fig. S1f and S1g), suggesting its crucial role in CRC initiation and disease progression.

Hence, we performed lentiviral shRNA-mediated stable silencing of *DKC1* in SW620, Colo320HSR, and WiDr cell lines. The *DKC1* knockdown was confirmed at both transcript and protein levels (Fig. S2a). Aligning with the role of dyskerin in cell cycle regulation, we also found a significantly reduced proliferation rate in *DKC1*-silenced CRC cells (**Fig. 2a**). Owing to the multifaceted role of DKC1 and its relatively unexplored potential in CRC pathogenesis, we performed transcriptome profile analysis using SW620 cells, *DKC1*-silenced (sh*DKC1*-1 and sh*DKC1*-2) and scrambled control (shSCRM) (Fig. S2b, Supplementary table 2a and 2b). Intriguingly, biological processes involved in cell cycle and negative regulation of proliferation were upregulated, whereas the processes related to cholesterol biosynthesis were downregulated which has been previously shown to promote stemness and drug resistance in CRC^18^ (**Fig. 2b**, Supplementary table 2c and 2d). Furthermore, Gene Set Enrichment Analysis (GSEA) revealed significant negative enrichment of G2/M cell cycle transition upon *DKC1* silencing (Fig. S2c). Next, to confirm the role of DKC1 in cell cycle regulation, propidium iodide-based flow cytometry was performed, and a notable increase in G2/M phase was recorded in *DKC1* knockdown cells (**Fig. 2c**). Although *DKC1* attenuation has been previously linked with G1/S mitotic checkpoint arrest and G2/M arrest in lung adenocarcinoma and sarcoma respectively^19, 20^, our findings here imply its context-dependent role in cell cycle progression. In agreement with this, we observed a significant decrease in the expression of genes involved in the G2/M phase transition (**Fig. 2d**).

**Figure 2:**
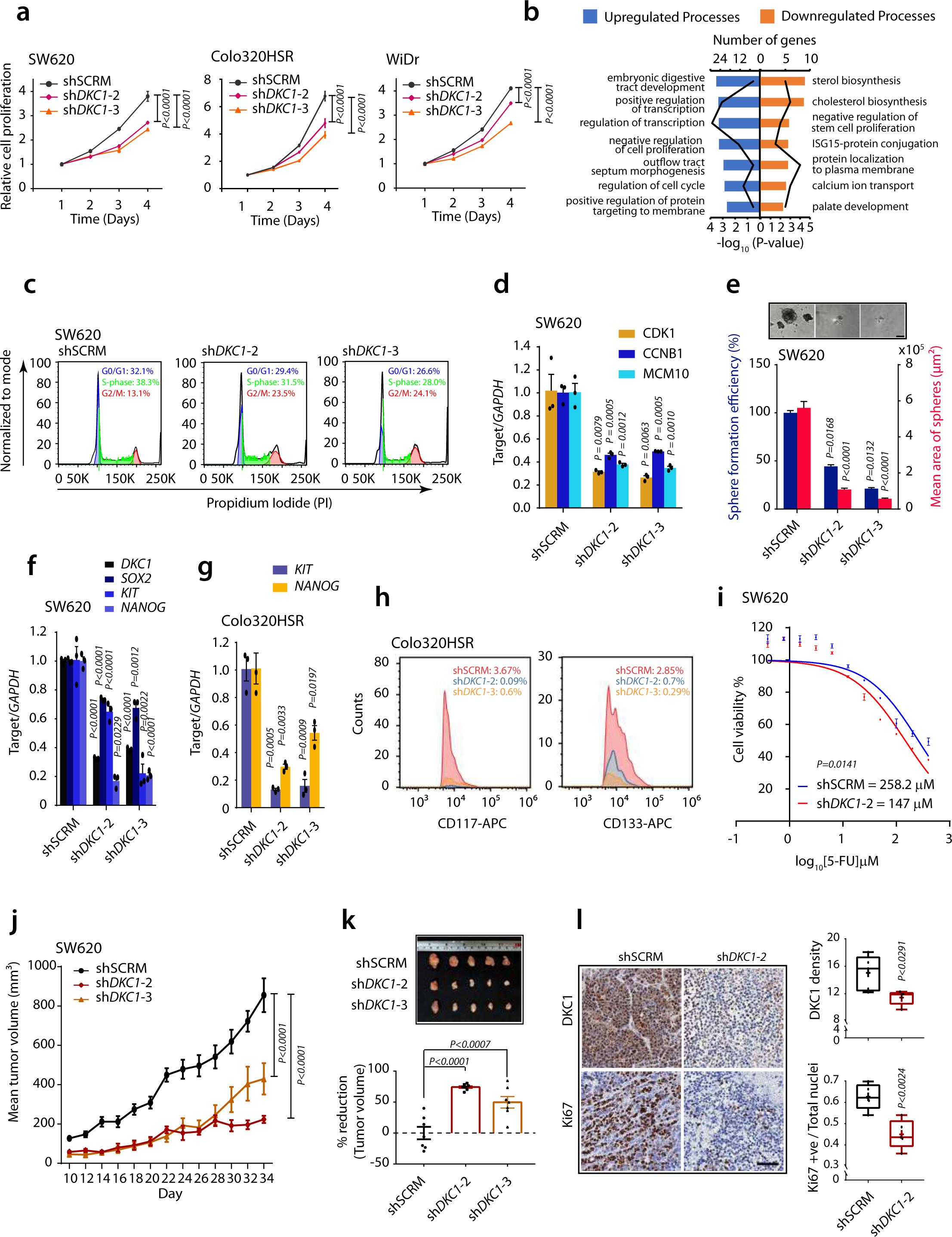
Attenuating *DKC1* in colorectal cancer reduces oncogenic properties and confers sensitivity to 5-fluorouracil. **a,** Cell proliferation using control (shSCRM) and *DKC1* knockdown (sh*DKC1*-2 and sh*DKC1*-3) SW620, Colo320HSR, and WiDr cells at the indicated time points. **b,** Upregulated and downregulated biological processes using RNAseq data of SW620 sh*DKC1* vs shSCRM cells by DAVID functional annotation clustering; bars (blue) represent −log10(P-value) and frequency polygon (red line) represent number of genes. **c,** Propidium iodide based cell cycle analysis for *DKC1* knockdown and control SW620 cells using flow cytometry. **d,** Expression of genes involved in cell cycle using same cells as in **c**. **e,** Tumor sphere formation assay with representative images (scale bar = 400μm) using same cells as in **c**; the blue bars represent sphere formation efficiency and the red bars show mean area of spheres. **f,** Relative expression of stemness genes in tumorspheres obtained from **e**. **g,** Relative expression of stemness markers in *DKC1* silenced and control Colo320HSR cells. **h,** Flow cytometry-based cell surface expression of CD117 and CD133 labeled with APC in same cells as in **g**, graphs represent percent positive cells. **i,** IC_50_ graphs for shSCRM and *DKC1* silenced SW620 cells using 5-fluorouracil. **j,** Mean tumor volume for control and *DKC1* silenced CRC cells xenografted in NOD/SCID mice. **k,** Bar graph depicting percentage reduction in tumor volume and representative image of tumors (upper panel). **l,** Immunostaining for DKC1 and Ki67 using control and *DKC1* knockdown tumors excised from xenografted mice (scale bar = 50μm). For panels, **a** and **j,** Two-way ANOVA, Dunnett’s multiple comparisons test was used for all the comparisons. **d, e, f, g,** and **l,** Unpaired Student’s two-tailed t-test was used. **k,** Unpaired Student’s two-tailed t-test with Welch’s correction were applied. Data are expressed as mean ± s.e.m.

Since DKC1 functions as a coactivator with SOX2/OCT4 complex and regulates the transcription of *NANOG* in embryonic stem cells^21^, we performed a sphere-formation assay using *DKC1*-silenced and control SW620 cells, and examined the ability of cancer stem cells to survive and form spheres in medium supplemented with minimal growth factors. A significant reduction in the sphere-forming efficiency and the mean surface area of the spheres was observed in *DKC1* silenced cells compared to control (**Fig. 2e**). Subsequently, the expression of stemness-associated markers and *DKC1* was examined in the tumor spheroids, and a significant reduction in the expression of *SOX2*, *KIT,* and *NANOG* was noted (**Fig. 2f**). Similar trend was observed in Colo320HSR cells, where a significant reduction in *KIT* and *NANOG* expression was observed upon *DKC1* silencing (**Fig. 2g**). Next, we examined the cell surface expression of CD117 (c-KIT) receptor tyrosine kinase, an established marker for colorectal cancer stem-like cells, and CD133 (Prominin I), a marker of stem cells or progenitor cells. Our flow cytometry data revealed a >6-fold decrease in the surface expression of CD117 and >4-fold reduction in CD133 expression in *DKC1* silenced Colo320HSR cells, a similar trend was observed for CD117 in *DKC1*-silenced SW620 cells, while no change was noted in CD133 (**Fig. 2h** and Fig. S2d). In line with this, a significant reduction in the foci formation ability of *DKC1-*silenced SW620 and WiDr cells was also observed (Fig. S2e and S2f). Next, to examine the role of DKC1 in drug resistance, we performed chemosensitivity assay, and an enhanced sensitivity of *DKC1* silenced cells to 5-fluorouracil was observed (**Fig. 2i**).

To investigate the role of DKC1 in tumorigenesis, we subcutaneously implanted the SW620- shSCRM and SW620-sh*DKC1* cells in NOD/SCID mice, and the tumor growth was monitored. A remarkable reduction (∼81%) in tumor burden was observed in the SW620-sh*DKC1* cells implanted group compared to the control group (**Fig. 2j and 2k**). Moreover, a significant decrease in the abundance of Ki67-positive nuclei, a marker for actively dividing cells, was observed in *DKC1*-silenced xenografted tumors compared to scrambled control (**Fig. 2l**). Taken together, we show that high DKC1 expression modulates oncogenic properties and drives colorectal tumorigenesis.

### DKC1 associates with DNA damage repair pathway

Since DNA damage response and checkpoint pathways often lead to cell cycle arrest^22^, we sought to explore if silencing *DKC1* in CRC induces DNA damage. We observed a strong positive correlation was found between DKC1 and genes involved in DNA repair machinery using publicly available CRC datasets (**Fig. 3a and 3b**). Next, we examined the phosphorylation of H2AX protein, γH2AX, a highly specific marker for double-strand DNA breaks, using *DKC1* silenced and control SW620 cells, and an increase in γH2AX positive foci was observed upon *DKC1* knockdown (**Fig. 3c**). Quantification of γH2AX revealed a significant increase in the number of foci per cell as well as in the frequency of γ-H2AX positive cells (**Fig. 3d**). Additionally, we checked the expression of DDR genes and a significant decrease in their expression upon *DKC1* silencing was observed (**Fig. 3e**). These results emphasize the role of DKC1 in maintaining genomic integrity in colorectal cancer.

**Figure 3:**
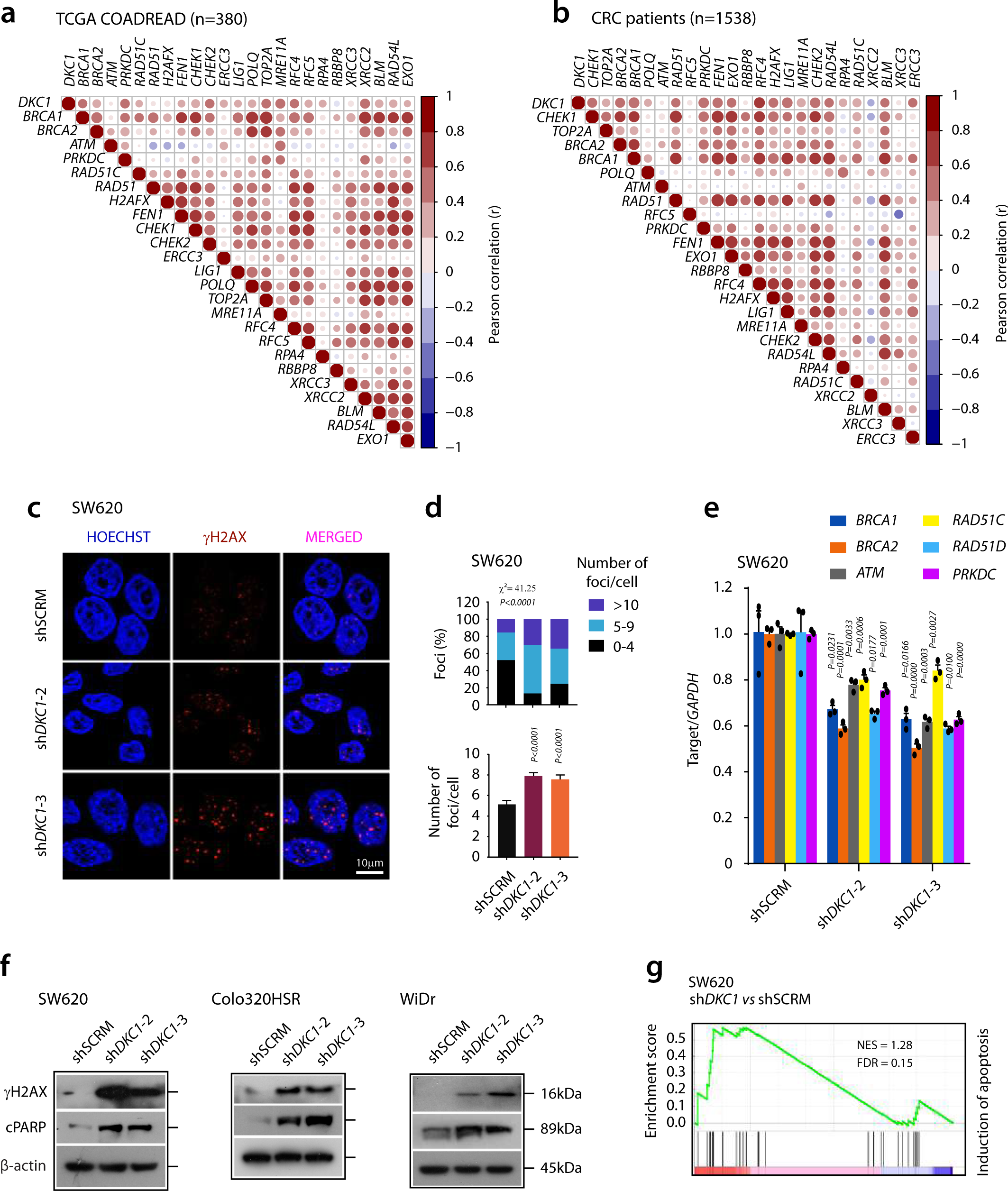
Silencing DKC1 in colorectal cancer triggers DNA damage and cell death. **a,** Correlation analysis of *DKC1* with DDR signature using data from TCGA COADREAD (N=380). **b,** Same as in **a** except multiple CRC datasets (N=1538) were used; scale bars represent Pearson’s correlation scores. **c,** Immunostaining for γ-H2AX using *DKC1* silenced and control SW620 cells; red puncta represent double-strand DNA breaks marked by γ-H2AX foci, Hoescht stain (blue) was used to mark nuclei (Scale bar = 10μm). **d,** Quantification of γ- H2AX immunostaining from **c**. **e,** Relative expression of DDR genes using same cells as in **c**. **f,** Immunoblots for markers for DNA damage and apoptosis using *DKC1* silenced and control SW620, Colo320HSR, and WiDr cell lines. **g,** GSEA plot showing enrichment of apoptosis using gene expression data for shDKC1 and control SW620 cells. For panel **d,** Unpaired Student’s two-tailed t-test with Welch’s correction (top panel) and Chi-square test for comparison of frequencies was used (bottom panel). **e,** Unpaired Student’s two-tailed t-test was applied. Data are expressed as mean ± s.e.m.

Poly ADP-ribose polymerase (PARP) is known to get recruited at the site of DNA damage for its repair, and the presence of cleaved PARP fragments indicates ongoing apoptosis^23^. Interestingly, we observed an increase in levels of cleaved-PARP in all three *DKC1-*silenced CRC cell lines accompanied by increased γH2AX levels (**Fig. 3f**). To check whether DNA damage is inducing apoptosis, we analyzed the transcriptome data for *DKC1* knockdown versus control cells, and a significant positive enrichment of pathways linked to apoptosis was observed (**Fig. 3g** and Fig. S2g). Collectively, our findings implicate a regulatory role of DKC1 in orchestrating DNA damage repair pathways.

### DKC1-positive patients associate with CMS2 class of colorectal cancer

Considering the highly heterogenous nature of CRC, we sought to examine whether the elevated DKC1 expression associates with a defined molecular signature or subtype. The widely accepted classification system for CRC is Consensus Molecular Subtypes (CMS), thus we employed the gene signatures for different CMS classes as CMS-UP or -DOWN (Supplementary table 3) and analyzed against the combined dataset of 770 CRC patients categorized as *DKC1*-high and *DKC1*-low groups. The Gene Set Variation Analysis (GSVA) shows upregulation of the ‘CMS2 UP’ gene signature in *DKC1*-high patients while a downregulation in *DKC1*-low patients (Fig. S3a). Next, we utilized ColoType, a 40 gene-signature for defining CMS class based on the global transcriptome data^24^, and found a similar association with ‘CMS2 UP’ gene signature (Fig. S3b). Subsequently, to validate this finding, we examined the publicly available transcriptome data for unsorted single cells (N=91,103) from two different CRC patient cohorts (GSE144735 and GSE132465)^25^, evaluated the average expression and percentage of cells expressing *DKC1* in different CMS subtypes. Both patient cohorts show a higher percentage of cells aligning to CMS2 subclass along with relatively high mean expression of *DKC1* compared to other subclasses (Fig. S3c and S3d). Further, a notable fraction of cells with high DKC1 expression exhibit Stem-like/TA (Transit Amplifying) phenotype in both the cohorts (Fig. S3c and S3d), which corroborates with our results where a remarkable reduction in stemness upon *DKC1* silencing was noted (**Fig. 2e**). Furthermore, we examined the association of differentially expressed genes between the comprehensive patient data (N=770) categorized based on DKC1 expression and the cell-type gene signature proposed in an earlier CRC classification^26^. A remarkable 14% overlap between the upregulated TA cells gene signature and upregulated genes in *DKC1*-high vs *DKC1*-low CRC patients was observed (Fig. S4A and Supplementary table 4). Furthermore, a notable ∼7% overlap of downregulated stem cell gene signature with downregulated genes in *DKC1*-high cases was noted. Next, we retrieve the proteomic profile of CRC patients from the Clinical Proteomic Tumor Analysis Consortium (CPTAC) and found an association of CRC patients with high DKC1 with CMS2 molecular subtype (**Fig. 4a**). Additionally, we examined the association of DKC1 with microsatellite and chromosomal instability phenotypes, and a significant association of high DKC1 with patients exhibiting microsatellite stable (MSS) and chromosomal instability-high (CIN-H) was observed (**Fig. 4b**). Collectively, we discover that CRC patients with high DKC1 strongly associate with CMS2 and TA/stem cell phenotypes.

**Figure 4:**
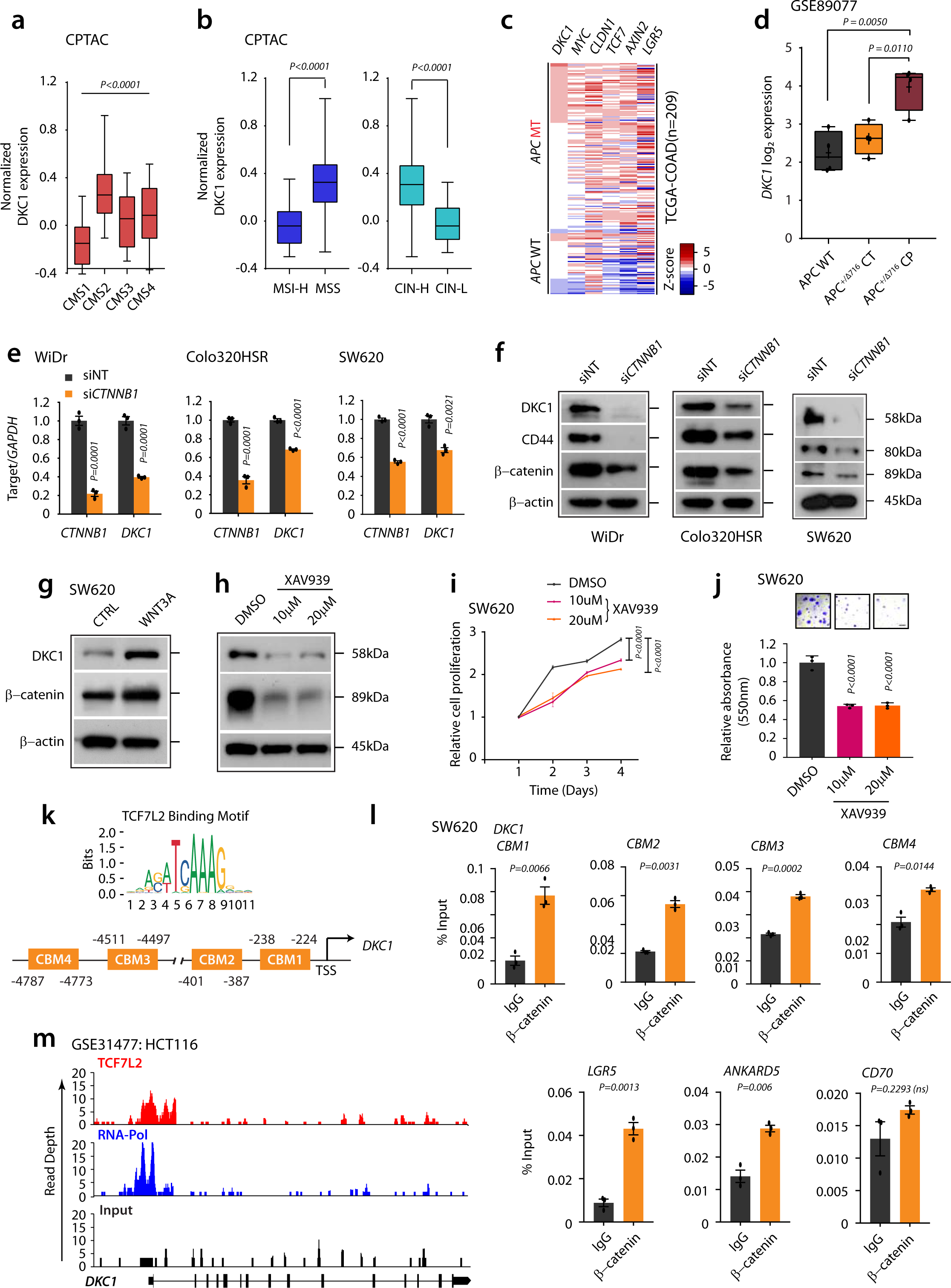
Canonical WNT signaling driven DKC1 associates with CMS2 subtype of CRC. **a,** Relative expression of DKC1 in CRC patient samples (N=76) categorised based on CMS classification; data retrieved from CPTAC. **b,** Relative expression of DKC1 in CRC patient samples (N=95) categorized based on microsatellite instability and chromosomal instability, data retrieved from CPTAC. **c,** Heatmap depicting expression of *DKC1* and WNT regulated genes in CRC patient specimens categorized based on *APC* mutation status. **d,** *DKC1* expression in colon tissue of APC wild type (APC WT), APC mutant (APC^+/Δ716^ CT) and colon polyps obtained from APC mutant (APC^+/Δ716^ CP) mice (GSE89077). **e,** Transcript levels of *DKC1* upon siRNA-mediated silencing of *CTNNB1* (β-catenin) in SW620, WiDr and Colo320HSR cells. **f,** same as in **e**, except protein expression, WNT target gene, CD44, was used as a positive control. **g,** Immunoblots showing DKC1 and β-catenin levels upon stimulation with conditioned media enriched with canonical WNT ligand, WNT3A. **h,** Immunoblots showing DKC1 and β-catenin levels in XAV939 treated SW620 cells. **i,** Cell proliferation using control and XAV939 treated SW620 cells. **j,** Foci formation assay using same cells as in **i** (scale bar = 1mm). **k,** Schema depicting the chromosomal location of β- catenin binding motif (CBM 1 to 4) at the *DKC1* promoter. **l,** ChIP-qPCR data showing enrichment of β-catenin upstream of *DKC1* using SW620 cells (*P* < 0.0001); *LGR5* and *ANKARD5* were used as positive controls and *CD70* as a negative control. **m,** ChIP-seq data showing enrichment of TCF7L2 and RNAPol binding on DKC1 promoter. For panels, **a,** One-way ANOVA, Dunnett’s multiple comparisons test was used. **b** and **d,** Unpaired Student’s two-tailed t-test with Welch’s correction was used. **e** and **l,** Unpaired Student’s two-tailed t-test was applied. **i,** Two way ANOVA, Dunnett’s multiple comparisons test was used. Data are expressed as mean ± s.e.m.

### Canonical WNT signaling drives *DKC1* in colorectal cancer

Since the CMS2 subclass of CRC invariably shows WNT signaling activation^8^, we sought to explore if *DKC1* expression is associated with active WNT signaling. Majority of CRC cases display activation of WNT signaling via inactivating mutation(s) in the *APC* tumor suppressor, thus using TCGA-COAD dataset the patients were stratified based on *APC* mutation and wildtype, and compared the expression of *DKC1* with other WNT target genes that play crucial roles in CRC tumorigenesis^27–31^. Interestingly, high *DKC1* expression exhibits a positive association with most of the *APC* mutant as well as wild-type cases accompanied with higher expression of WNT targets (**Fig. 4c**). In accordance, *DKC1* shows a positive correlation with WNT signaling effector *CTNNB1* in TCGA COADREAD dataset (Fig. S4b). Additionally, colorectal polyps from *APC*^+/Δ716^ mice show significantly high *DKC1* expression compared to normal colorectal tissues obtained from *APC*^+/+^ and *APC*^+/Δ716^ mice (**Fig. 4d**, GSE89077). To confirm the direct control of WNT signaling on the *DKC1* expression, we analyzed a publicly available dataset (GSE44097), and a significant reduction in the *DKC1* expression upon *CTNNB1* silencing in CRC cells was obtained (Fig. S4c). Moreover, siRNA-mediated silencing of *CTNNB1* in WiDr, Colo320HSR, and SW620 cells led to a significant decrease in DKC1 expression, and CD44, a target of WNT signaling was used as a positive control (**Fig. 4e and 4f**). In line with this, stimulation of SW620 cells with canonical WNT ligand WNT3A significantly increased DKC1 expression, whereas tankyrase inhibitor XAV939 markedly reduced DKC1 expression (**Fig. 4g and 4h**). Additionally, treating SW620 cells with XAV939 reduced proliferation rate and foci formation ability of these cells proving effectiveness of WNT signaling inhibitor against high DKC1 expressing CRC (**Fig. 4i and 4j**). Further, to prove WNT signaling mediated upregulation of *DKC1*, we examined its upstream region for the presence of TCF/LEF binding motifs using the JASPAR CORE transcription factor binding site prediction tool^32^. We found several putative TCF7L2 binding sites or Catenin Binding Motifs (indicated as CBM1-CBM4) in the proximal and distal promoters of *DKC1* (**Fig. 4k**). To confirm the occupancy of β-catenin/TCF complex, we performed chromatin immunoprecipitation (ChIP)-PCR using β-catenin antibody using SW620 cells; as expected, a significant enrichment of β-catenin/TCF complex at all the four CBMs was observed (**Fig. 4l**). Further, in conjunction with our finding, a publicly available ChIP-Seq dataset (GSE31477) revealed enrichment of TCF7L2 binding along with RNA polymerase II in HCT116 cells, indicating the active transcriptional status of *DKC1* (**Fig. 4m**). Taken together, our findings established direct control of canonical WNT signaling on the transcriptional regulation of *DKC1*.

### DKC1 plays a crucial role in sphingolipid biosynthesis

Lipid metabolism provides a targetable metabolic vulnerability in CRC, and the clinical utility of lipid-based signatures received much attention as a non-invasive diagnostic and prognostic biomarker^33^. The GSEA of our transcriptomics data indicates the downregulation of primary biological processes responsible for lipid production upon silencing *DKC1*, such as lipid and Acetyl Co-A metabolism (**Fig. 5a** and Fig. S5a). Furthermore, we performed lipid profiling of *DKC1* silenced and control SW620 cells using ultra-high performance liquid chromatography coupled with tandem mass spectrometry (UHPLC-MS/MS) and employed multivariate analysis. The principal component analysis revealed two distinct clusters highlighting the varied lipidome of *DKC1* silenced versus control cells (**Fig. 5b**). Intriguingly, the differential lipid profile revealed significant downregulation of various ceramides and sphingosine, while a concurrent upregulation of several phospholipids and sphingomyelins were observed in *DKC1*-silenced cells compared to control (**Fig. 5c**, Supplementary table 5a). Next, we sought to identify the most important deregulated lipid moieties, and several phospholipid species including phosphatidylcholines, phosphatidylethanolamines, and various plasmalogen isoforms were found to be upregulated whereas several ceramide species including Gb3(d18:1/16:0), HexCer(d18:1/16:0) and Hex2Cer(d18:1/24:1) were downregulated with a VIP score of >1 (**Fig. 5d**, Supplementary table 5b). Furthermore, the Lipid Ontology network analysis (LION) revealed sphingolipid metabolism-related ontologies to be most significantly downregulated upon *DKC1* silencing (**Fig. 5e**, Supplementary table 5c), while glycerophospholipids related ontologies were upregulated. Nonetheless, the GSEA exhibits a negative enrichment of sphingosine biosynthesis in *DKC1*-silenced cells compared to scrambled control (Fig. S5b). Collectively, our findings highlight the significant role of DKC1 in lipid metabolism in CRC pathogenesis.

**Figure 5:**
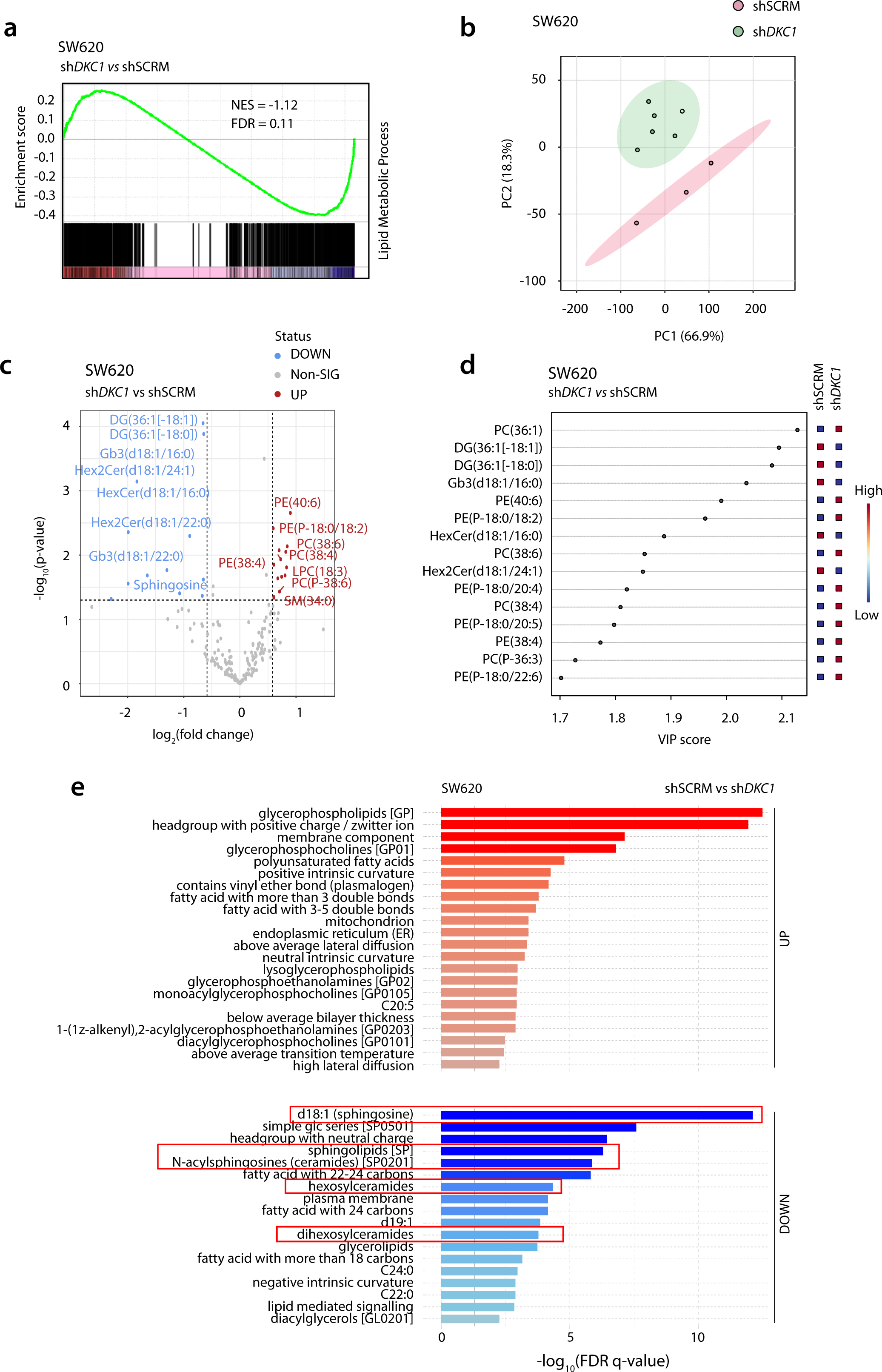
*DKC1* perturbs lipid metabolism in colorectal cancer. **a,** Gene set enrichment analysis for lipid metabolic biological process using RNAseq data of the*DKC1* knockdown and control SW620 cells. **b,** Principal component analysis of the lipidomics data using same cells as in **a**; unsupervised PCA defines the two groups invariably (PC1 and PC2 are Principal Component 1and Principal Component 2, respectively). **c,** Volcano plot depicting various lipid species showing more than or less than 1.5-fold up or down regulated levels upon DKC1 silencing in SW620 cells (dashed horizontal line represents P value of 0.05). **d,** Variable importance of projection (VIP) score plot shows important features (VIP score > 1.7). **e,** Lipid ontology (LION) network analysis using *DKC1* silenced and control SW620 cells; red bars depict upregulated ontologies whereas blue bars depict downregulated ontologies; significantly downregulated sphingolipid metabolism related ontologies are marked in red boxes.

### CRC patients with high DKC1 show an accumulation of ceramides

To investigate the DKC1-mediated dysregulation of lipid metabolism in CRC patients, DKC1 levels were analyzed in the tumor specimens and paired adjacent normal tissues (N=9), categorizing them as DKC1-high or DKC1-low tumors and subjected to lipidome analyses (Fig. S5C). The lipid profile revealed increased levels of Cer(d18:2/23:0) and Cer(d18:1/24:0) in *DKC1-*high cases, accompanied by low sphingomyelin (**Fig. 6a**, Supplementary table 6a). We next identified the most important variables in the differential lipid profile; wherein both the ceramides had VIP scores of >1.7, highlighting their importance in the prediction of *DKC1*- high cases (**Fig. 6b**, Supplementary table 6b). We next checked the efficacy of these two ceramides as biomarkers for *DKC1* positive cases and found a higher accumulation of the same in *DKC1-*high tumors, representing an area under the receiver operating characteristic (AUROC) curve of >0.9 showing high sensitivity and specificity (**Fig. 6c and 6d**). To further confirm, ROC curve-based model evaluation was performed using the Biomarker analysis module of Metaboanalyst 5.0 using the partial least square discriminant analysis (PLS-DA) for performance evaluation. An AUROC curve of 0.855 was noted for these ceramides, signifying their utility as biomarkers for differentiating *DKC1*-high vs. *DKC1*-low CRC (**Fig. 6e**). Furthermore, the Lipid Ontology Network analysis revealed upregulation of ontologies related to sphingosine and ceramides accompanied with downregulation of sphingomyelin related ontology in *DKC1*-high tumors, signifying the association of DKC1 with ceramide accumulation (**Fig. 6f**, Supplementary table 6c). Hence, we conclude that the sphingolipid metabolism is influenced by DKC1, and it plays crucial role in the modulation of ceramide levels in CRC patients.

**Figure 6:**
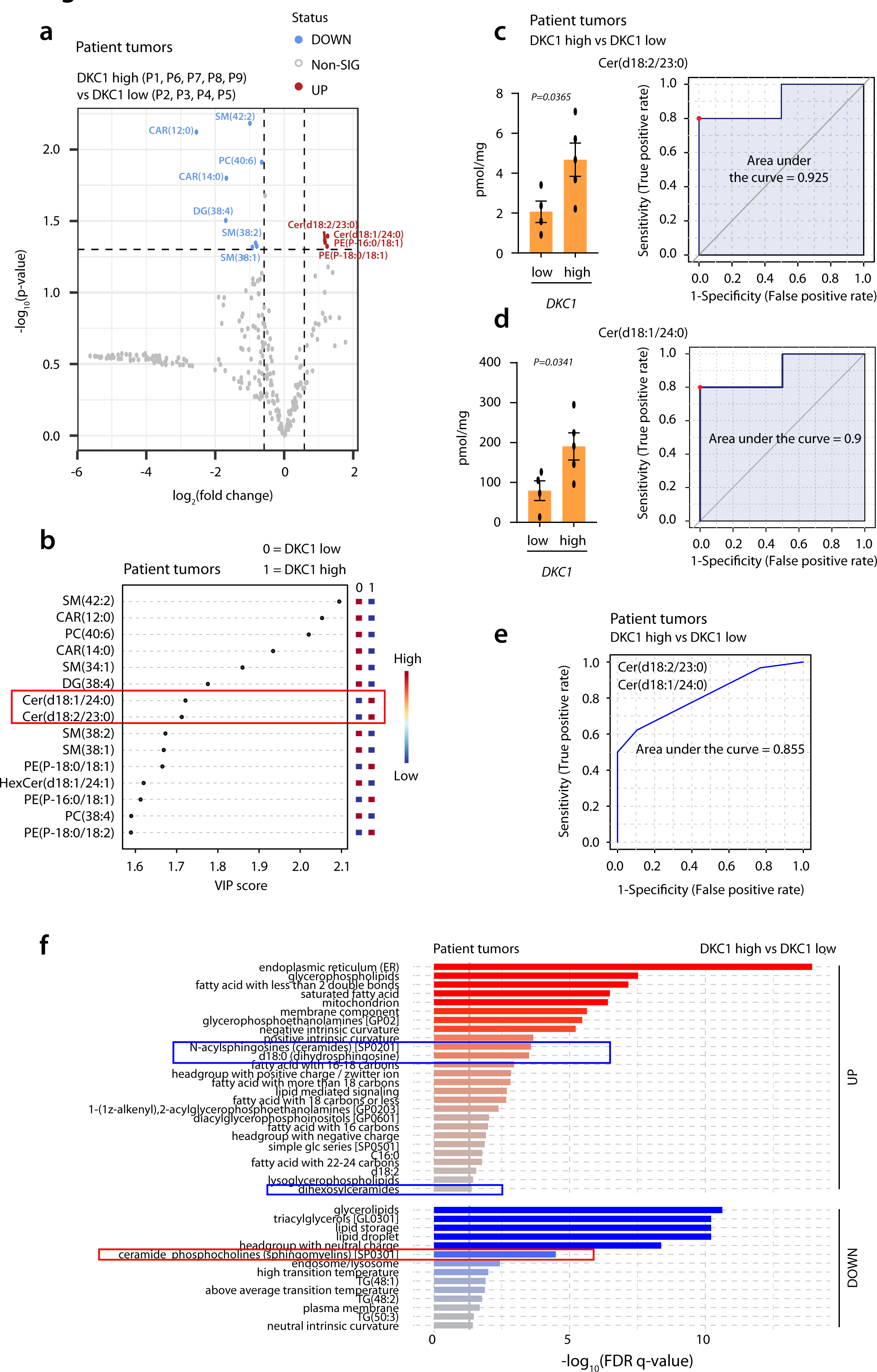
High DKC1 levels associate with upregulated sphingolipid biosynthesis in colorectal cancer patients. **a,** Volcano plot depicting lipid species more than or less than 1.5- fold up or down regulated in *DKC1*-high (P1T, P6T, P7T, P8T and P9T) and *DKC1*-low (P2T, P3T, P4T, P5T) CRC patient specimens, (dashed horizontal line represents P-value of 0.05). **b,** Variable importance of projection (VIP) score plot shows important features (VIP score > 1.5); C23 and C24 ceramide species were upregulated in *DKC1*-high tumors (marked in red box). **c,** Concentration of Cer(d18:2/23:0) using *DKC1*-high and *DKC1-*low tumor specimens and receiver operating characteristic (ROC) curve applying univariate analysis. **d,** Same as in C, except for Cer(d18:1/24:0). **e,** ROC curve obtained using Partial Least Square Discriminant Analysis based model evaluation for Cer(d18:2/23:0) and Cer(d18:1/24:0). **f,** Lipid ontology network (LION) analysis using *DKC1-*high and *DKC1* low CRC specimenss; red bars depict upregulated ontologies and blue depict downregulated ontologies; sphingolipids and ceramides related ontologies are significantly upregulated (marked in blue box) while sphingomyelin related ontologies are downregulated (marked in red box) in *DKC1*-high tumors.

### DKC1-driven *SGPP2* regulates sphingolipid biosynthesis

To interrogate the underlying regulatory mechanism involved in altered lipid metabolism in the context of DKC1*-*mediated CRC, we performed integrative pathway analysis using transcriptome and lipid profiling data of *DKC1*-silenced and control cells. In line with our patient lipidomics data, sphingolipid metabolism was noted to be most significantly affected with a high pathway impact score (**Fig. 7a**). Additionally, several metabolic hits such as sphingosine, lactosylceramide, digalactosylceramide and sphingomyelin were observed in the sphingolipid metabolism, while a single gene hit, namely *SGPP2* was recorded from the expression data (Fig. S5d).

**Figure 7:**
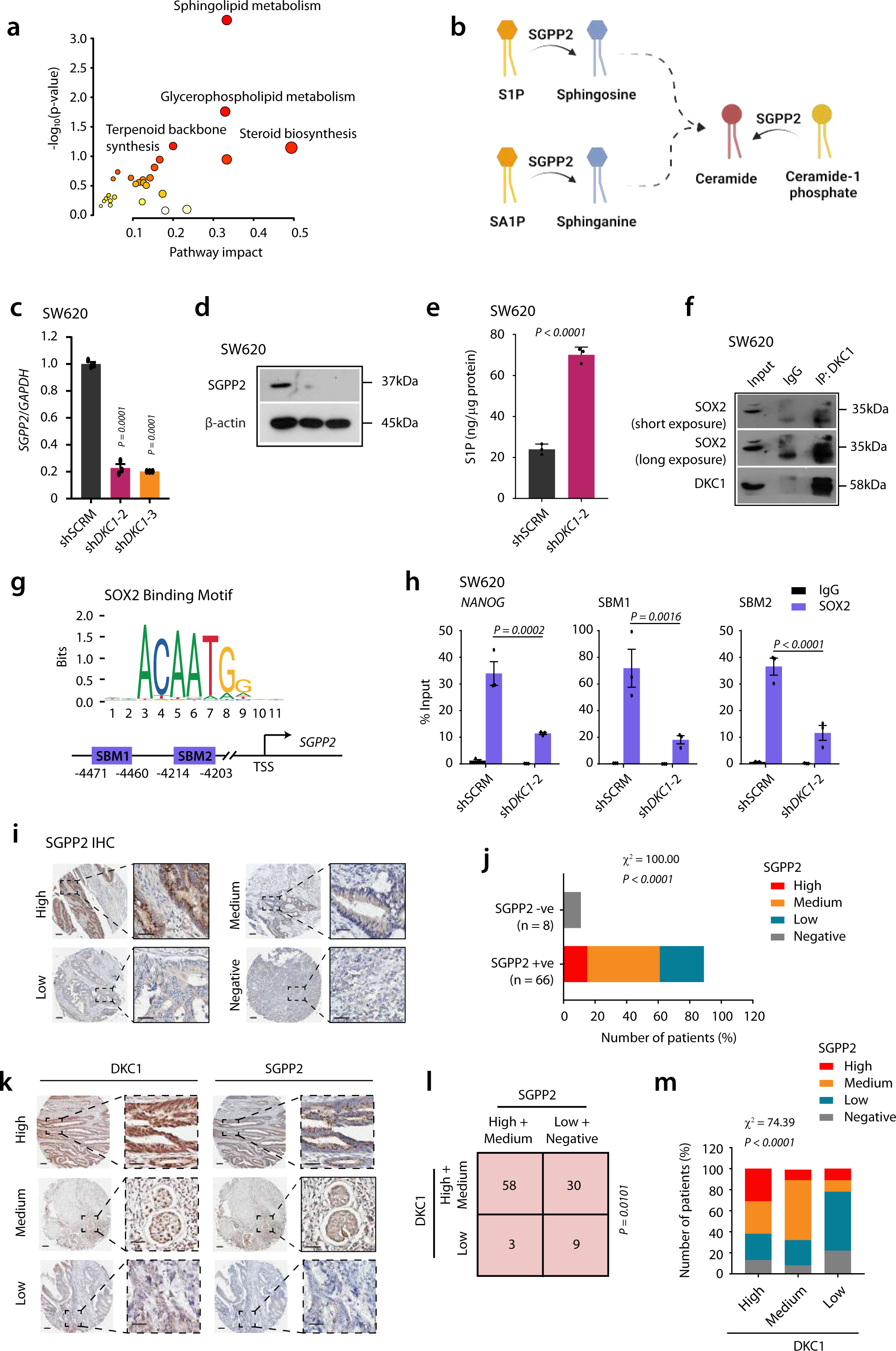
DKC1-mediated upregulation of *SGPP2* modulates sphingolipid metabolism in colorectal cancer. **a,** Integrative analysis of the RNAseq and lipidomics datausing DKC1 silenced and control SW620 cells, -log_10_(P-value) of >1.3 was considered significant. **b,** Schema showing SGPP2-mediated dephosphorylation of sphingosine-1-phosphate (S1P), sphinganine-1-phosphate (SA1P) and ceramide-1-phosphate, leading to accumulation of ceramides. **c** and **d,** Transcript and protein levels of SGPP2 upon *DKC1* silencing in SW620 cells. **e,** S1P levels in the DKC1 knockdown and control SW620 cells. **f,** Immunoprecipitation data showing interaction between SOX2 and DKC1 using SW620 cells. **g,** Schema showing predicted SOX2 binding sites on the *SGPP2* promoter (SOX2 binding motifs 1 and 2 marked as SBM1 and SBM2). **h,** ChIP-PCR data for SOX2 binding on the *SGPP2* promoter using *DKC1* knockdown and control SW620 cells. **i** and **j,** Representative immunostained images of CRC patient specimens (N=74) for SGPP2 expression,; ∼90% of the CRC cases show SGPP2 expression as depicted in bar plot. **k,** Representative immunostained images for expression of DKC1 and SGPP2 in CRC patient specimens. **l,** Contingency analysis of DKC1 and SGPP2 expression for CRC patients. **m,** Bar plot showing co-expression of DKC1 and SGPP2 in CRC patient specimens. For panel **c, e** and **h,** Unpaired Student’s two-tailed t-test was used. **j** and **m,** Chi square test for comparison of frequencies was used for both panels. **l**, Fisher’s exact test for contingency analysis was used. For **i** and **k,** scale bar for whole section is 100 μm and for inset image is 40 μm. Data are expressed as mean ± s.e.m.

Sphingosine-1-phosphate phosphatase 2 (*SGPP2*) catalyzes the dephosphorylation of sphingosine-1-phosphate (S1P) to sphingosine, sphinganine-1-phosphate to sphinganine, and ceramide-1-phosphate to ceramide thereby regulating the production of ceramides (**Fig. 7b**). Thus, we next examined the expression of *SGPP2* in DKC1 knockdown cells, and a significant reduction in SGPP2 levels was observed, which indeed aligns with our lipid profiling data wherein a significant reduction in sphingosine and ceramide levels was noted (**Fig. 7c and 7d**). Furthermore, to examine whether reduced *SGPP2* expression upon *DKC1* silencing is hampering the conversion of S1P to sphingosine, we analyzed S1P levels upon DKC1 silencing, an increased accumulation of S1P was observed in *DKC1* knockdown cells compared to control SW620 cells (**Fig. 7e**), indicating the key role of DKC1 in the regulation of *SGGP2* and sphingosine metabolism. Since DKC1 forms a complex with SOX2/OCT4 and regulates the expression of *NANOG* in embryonic stem cells ^21^, we conjectured regulation for *SGPP2* via this transcriptional complex. To confirm this interaction, immunocomplex was pulldown using DKC1 antibody and immunoblotted for SOX2, a notable expression of SOX2 confirms its interaction with DKC1 (**Fig. 7f**). Furthermore, the regulatory region of *SGPP2* was scanned using JASPAR CORE transcription factor binding site prediction tool, and two distinct SOX2 binding motifs (SBM1 and SBM2) within 5 Kb upstream to the transcription start site (TSS) were identified (**Fig. 7g**). To examine the binding of DKC1/SOX2 complex on the promoter of *SGPP2*, chromatin immunoprecipitation-PCR (ChIP-PCR) was performed and a remarkable enrichment of SOX2 at both the SBMs in the control cells was observed, while significantly reduced enrichment was noted in *DKC1* silenced cells (**Fig. 7h**). Taken together, we established DKC1 – SOX2 complex mediated regulation of *SGPP2,* and its significance in ceramide accumulation in CRC.

To cross-examine the regulatory interplay between DKC1 and SPGG2 in CRC patients, the tissue microarray sections comprising tumor specimens (N=74) of varying grades and stages were immunostained for DKC1 and SGPP2 expression. In concordance with the role of DKC1 in transcriptional regulation of *SGPP2,* its expression is ubiquitously present primarily in the nucleus, and to some extent in the cytosol in almost all patient specimens. Albeit owing to the enzymatic function of SGPP2, its expression was predominantly cytoplasmic. Majority of the CRC tumors (∼89%) show positive staining for the SGPP2 expression (**Fig. 7i and 7j**), wherein ∼15% exhibit high SGPP2 levels, moderate in ∼46%, and low in ∼28% of the patients, while ∼11% show negative staining for SGPP2. About ∼58% of patients show high to medium expression of both SGPP2 and DKC1, while ∼30% of cases (DKC1-high and medium) exhibit low to negative expression of SGPP2 (**Fig. 7k and 7l**). Furthermore, ∼62% of the DKC1-high cases show either high or medium expression of SGPP2, while most DKC1 low patients harbor either low or negative SGPP2 expression (**Fig. 7m**). Conclusively, we deciphered the molecular mechanism involved in the accumulation of ceramides in DKC1-positive tumors, signifying the importance of these lipid-based biomarkers in further defining the molecular subtypes of CRC.

## DISCUSSION

In this study, we deciphered the functional significance of DKC1 as an oncogene, show its association with disease recurrence, and discovered its transcriptional regulation via canonical WNT signaling. Nonetheless, DKC1 positive CRC exhibits remarkable overlap in the molecular signatures including WNT targets associated with the CMS2 class. In congruence, our results demonstrate the utility of WNT inhibitors in abrogating the DKC1-mediated oncogenic properties. Importantly, in this study, we discovered disturbed metabolism of lipids leading to the accumulation of ceramides through DKC1-mediated upregulation of SGPP2. Thus, we propose the importance of DKC1-mediated deregulation of sphingolipid metabolism and highlight the clinical utility of ceramides as biomarkers for the identification of DKC1- positive cases (**Fig. 8**).

**Figure 8:**
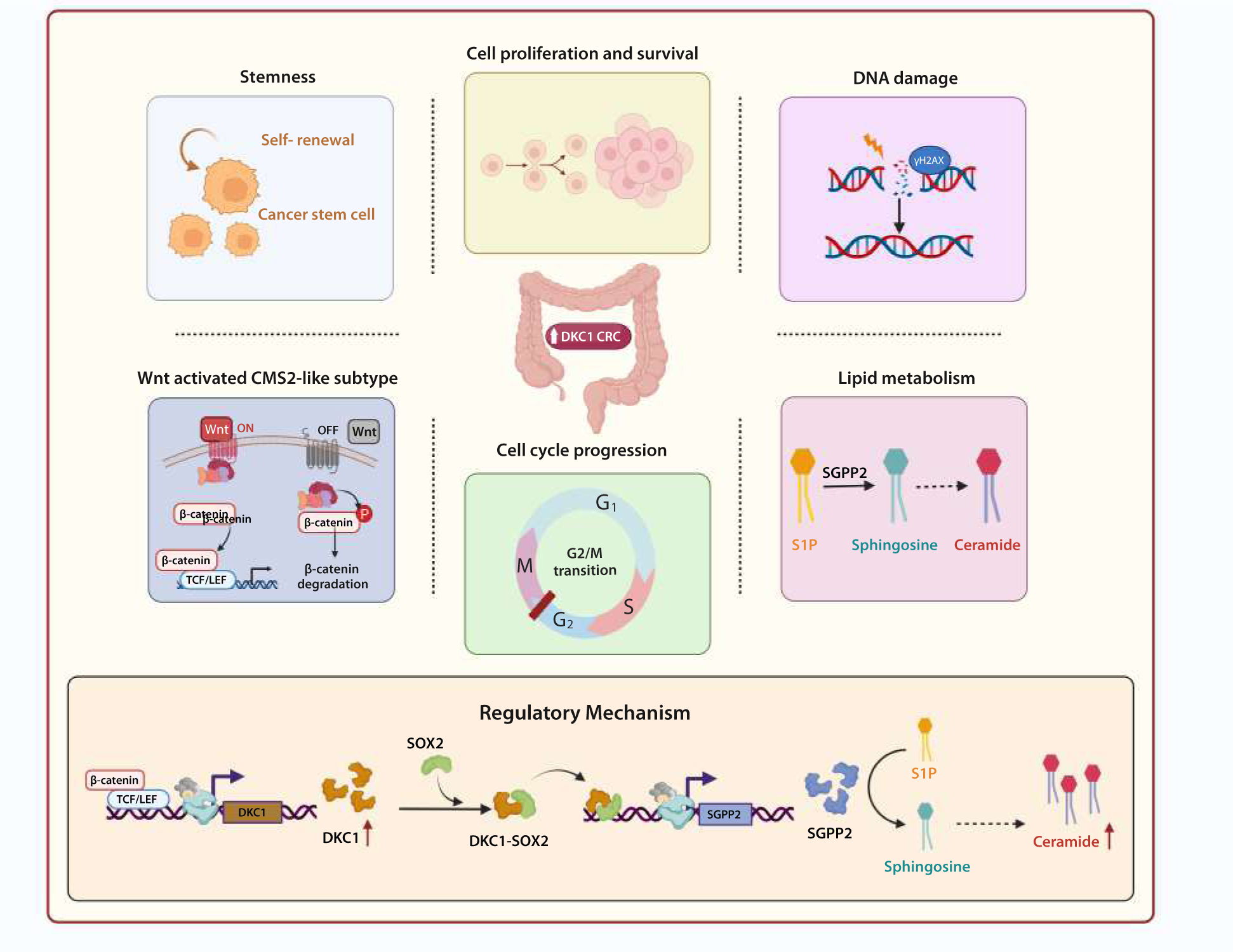
Schematic overview showing oncogenic roles mediated by DKC1 in colorectal cancer progression.

Previously, impaired DKC1 activity has been linked with cell cycle arrest in dyskeratosis congenita, wherein defective ribosome often leads to bone marrow failure, skin abnormalities, and increased susceptibility to cancer^36^. In accordance, inactivating mutation in *DKC1* has been linked with reduced expression of p53 via defective IRES-mediated mRNA translation in breast cancer^38^. In line with this, our findings affirm the role of DKC1 in DNA double-strand break repair and apoptosis. The high ploidy and chromosomal instability in cancer have been attributed to defects in homologous recombination (HR) and non-homologous end joining (NHEJ) pathways^39, 40^. Moreover, the CMS2 class has also been linked to high ploidy with activation of the HR pathway^41^. Notably, a key player in NHEJ, namely, PRKDC, has been defined as an oncogene in CRC^42^, which in our study was found to be downregulated upon *DKC1* silencing.

Distinct CMS classes require specific delineation of molecular and phenotypic features for the identification of novel drug targets. Moreover, CMS2 exhibits poor immunogenic signature, and most of the patients belonging to this class show an association with tubular adenomas and epithelial signature. Molecularly, this specific class harbour TA cell signature, abundance of *APC* and *TP53* mutations, low methylation pattern, and enhanced ribosomal translation^43^. Most of these features aligns with ours and can be traced back to decide evidence-based effective therapeutics.

The FOLFIRI-based treatment has been more effective against stemlike phenotype in cell type-based CRC classification. While the transit amplifying class show differential response to cetuximab and was further divided into two subclasses, cetuximab sensitive and resistant ^26^. In this study, *DKC1* associate with stemness features and its expression in single-cell data analysis further confirms stemlike/transit amplifying phenotype. Thus, targeting *DKC1* using WNT inhibitors in cetuximab-resistant transit amplifying class could provide a better alternative treatment avenue for RAS/RAF mutant CRC patients.

Ceramide accumulation has been shown to inhibit mitochondrial fatty acid oxidation^46^; likewise in this study, DKC1-mediated dysregulation of lipid metabolism which led to higher levels of ceramides is speculated to reduce fatty acid oxidation. The decrease in ceramides could be dependent on the reduced sphingosine levels which generates ceramides through the salvage pathway^47^. Furthermore, in line with the known transcriptional co-activator activity of DKC1^21^, its regulatory role in *SGPP2* expression provided novel concrete evidence of the direct involvement of DKC1 in sphingolipid metabolism. Complex sphingolipids have been known to play a crucial role in tumorigenesis^48^, thus, the presence of lactosylceramide and galactosylceramides in our integrative analysis indicates pro-survival roles for these lipids. Moreever, regulation of glycosylated ceramide and sphingomyelin efflux is crucial for imparting drug resistance^49, 50^, therefore, their plausible role needs to be explored in the context of DKC1-positive CRC. Accumulation of C24 ceramide in DKC1-positive patients as shown in this study, is in line with its known role of partial prevention from programmed cell death^51^.

Since, the biomarker identification for defining molecular subsets in a cancer offer further development of diagnostics or therapeutic approaches, our discovery of DKC1 positive subset-specific ceramide biomarkers require validation in a larger CRC cohort. Though we do not neglect the probability of finding other ceramide species, DKC1-mediated dysregulation of sphingolipid metabolism may offer promising biomarkers.

Collectively, we highlighted the poor prognosis associated with high *DKC1* expression and established the crucial role of the WNT-DKC1 axis in CRC progression, specifically in patients belonging to canonical CMS2 class, thus inhibitors against the WNT signaling pathway would be effective for these patients. Additionally, the involvement of DKC1 in the transcriptional regulation of sphingolipid metabolism and ceramide accumulation will open new avenues to be explored in CRC pathogenesis and therapeutics. Importantly, methods implying the detection of ceramide species as a biomarker(s) will aid in improvised diagnostic approaches for *DKC1*-positive colorectal cancer.

## MATERIALS AND METHODS

### Human CRC patient specimens

Surgically resected tumors and paired adjacent normal tissue samples were collected from Regency Hospital (Kanpur, India) or King George’s Medical University (Lucknow, India). The samples were collected after obtaining informed consent and institutional ethics board approvals as per the guidelines of the Indian Council of Medical Research (ICMR). All patients’ specimens used in this study were collected in accordance with the Declaration of Helsinki. A total of nine pair of fresh samples were collected and snap frozen using liquid nitrogen for preserving the metabolic caricature of the samples. Patient tissue microarray (TMA) slides (CO1502) comprising colorectal cancer specimens (N=74) with two cores for each patient were procured from Tissuearray.com LLC.

### Survival analysis

To check the association of DKC1 expression with the overall survival of colorectal cancer patients The Cancer Genome Atlas- Colon Adenocarcinoma (TCGA-COAD) dataset was downloaded from the UCSC Xena browser (https://xenabrowser.net). R packages Survival, Survminer, and SurvMisc were used for survival data analysis, cutoff optimization was done according to Cox proportional hazard regression model, and making Kaplan-Meier plot. Statistical significance was measured using log-rank test.

### Differential expression analysis of patient transcriptome data

Publicly available gene expression data for primary tumor samples (n = 1644) in colorectal cancer microarray datasets (GSE2109, GSE13067, GSE13294, GSE14333, GSE17536, GSE20916, GSE33113, GSE35896 and GSE39582) from Gene Expression Omnibus (GEO) were subjected to normalization using single sample frozen RMA method ^53^ from ‘frma’ package R/Bioconductor. These datasets were next subjected to outlier sample detection for quality assessment using ‘arrayQualityMetrics’ package from R ^54^. The outliers were excluded based on MA plots, pairwise sample distances, and gene expression value distribution. The Median Absolute Deviation (MAD) score was calculated for every probe in the remaining samples (n = 1538). While converting probes to gene symbols, duplicate symbols were removed by keeping the probe with the highest MAD score and rejecting others for each gene. Next, samples were subjected to quartile classification based on the expression of *DKC1* and the first and fourth quartiles were designated as *DKC1*-high and *DKC1*-low cases (n = 770). ‘limma’ package from R/Bioconductor was used to identify differentially expressed genes between *DKC1*-high and *DKC1*-low patients. Genes with log_2_(fold change) >= 1 were designated as upregulated and genes with log_2_(fold change) <= -1 were called as downregulated. Genes showing a positive association with *DKC1* expression were subjected to functional annotation clustering analysis using The Database for Annotation, Visualization, and Integrated Discovery (DAVID) ^55^ and Gene Ontology analysis was performed to analyze upregulated biological processes and pathways.

### CCLE data analysis

Publicly available expression data for *DKC1* was downloaded from from the DEPMAP database Cancer Cell Line Encyclopedia (https://depmap.org/portal/). Chronos and CERES scores for DKC1 were also retrieved from DEPMAP and plotted in GraphPad Prism 7.0.

### Cell culture and *DKC1* knockdown

Colo320HSR cells were obtained from American type culture collection (ATCC) and WiDr cells were obtained as a generous gift from Dr. Eric Fearon’s laboratory at the University of Michigan, Ann Arbor, MI, USA. SW620 cells were obtained from the Cell line repository at National Centre for Cell Sciences, Pune, Maharashtra, India. All three cell lines were validated by STR profiling (DNA Forensics Laboratory, New Delhi, and DNA Labs India, Hyderabad, India) and were maintained as per ATCC recommendations. Colo320HSR cells were maintained in RPMI1640 media whereas WiDr and SW620 cells were maintained in MEM and Leibovitz’s L-15 medium respectively, supplemented with 10% FBS and 0.5% Penicillin/Streptomycin. All cell lines were routinely tested for Mycoplasma contamination using the PlasmoTest mycoplasma detection kit (InvivoGen). For stable silencing of *DKC1*, human lentiviral short-hairpin RNA-targeting *DKC1* (sh*DKC1*) and non-targeting scrambled short-hairpin RNA (shSCRM) cloned in pLV vectors (Cyagen Biosciences, Santa Clara, USA) and packaged using ViraPower Lentiviral Expression System (Invitrogen, Carlsbad, CA, USA) as per manufacturer’s instructions.

### Real-time quantitative PCR

Real time expression of genes was analyzed using quantitative PCR as per the established protocol ^56, 57^. Briefly, RNA was isolated from the colorectal cancer cells using TRIzol reagent (Ambion, Life technologies) as per the instructions provided by the manufacturer, and 1μg of RNA was converted to cDNA using Random primers. qPCR was performed using SYBR Green PCR mastermix (Genetix Biotech Asia Pvt. Ltd) on the QuantStudio Real-Time PCR System (Thermo Fisher Scientific). Primer sequences are provided in the supplementary table 7.

### RNA sequencing and analysis

RNA was isolated as described above and exported to Macrogen Inc. (South Korea). Quality of RNA was assessed using 2100 Bioanalyzer (Agilent Technologies) followed by library preparation (using Illumina TruSeq Stranded mRNA Library method) and paired-end sequencing on NovaSeq with a 150bp readlength and more than 25M reads were recorded for each sample. Samples were subjected to differential expression analysis using DESeq2 ^58^ to identify dysregulated genes (> or < 0.6 log_2_(fold-change) and adjusted p-value <0.05). This gene list was subjected to functional annotation clustering-based Gene Ontology analysis using The Database for Annotation, Visualization, and Integrated Discovery (DAVID) ^55^.

For performing the Gene Set Enrichment Analysis (GSEA), samples were normalized and the batch correction was performed using Combat in R. Genesets with Normalized Enrichment Score (NES) of >1 or <-1 with a False Discovery Rate (FDR) of <0.25 were considered significant with GSEA being run with signal-to-noise-ratio as per the default parameters.

### Immunoblotting

Immunoblotting was performed as per the established protocol ^59^. Briefly, radioimmunoprecipitation assay (RIPA) buffer with Phosphatase Inhibitor Cocktail Set-II (Calbiochem) and protease inhibitor (VWR) was used to prepare whole cell protein lysates. Protein quantification was performed using BCA and proteins was separated on SDS-PAGE followed by transfer to PVDF membrane (GE Healthcare). Next, the membrane was incubated with 5% non-fat dry milk in Tris buffered saline with 0.1% Tween-20 (TBS-T) for an hour at room temperature. Antibody exposure was provided at the respective dilutions in TBS-T containing 2% non-fat dry milk: 1:1000 diluted DKC1 (CST, 53234), cPARP (CST, 9541), γ- H2AX (CST, 9718), SOX2 (Abcam, ab97959), 1:3000 diluted β-catenin (CST, D10A8) and 1:5000 diluted β-actin (Abcam, ab6276) followed by overnight incubation at 4°C. Post incubation, the antibody was removed, and the membrane was washed with 0.1%TBS-T and was incubated for 2hr at room temperature with respective secondary anti-mouse or anti-rabbit antibody conjugated with horseradish peroxidase (HRP) (1:5000, Jackson ImmunoResearch Laboratories, Cat # 115-035-003 and Cat # 111-035-144). Again, the membrane was washed with 0.1% TBS-T and the signal was captured using an enhanced chemiluminescence system (ThermoFisher).

### Cell proliferation assay

Cells were trypsinized and counted using Countess 3 cell counting system (Invitrogen) and 2×10^3^ cells/per well were plated in a 96 well plate for SW620 and WiDr cells (1×10^3^ cells were plated for Colo320HSR cells). Resazurin was added to the cells and fluorescence intensity was measured at respective time points at 530-590 nm excitation-emission. Relative cell proliferation rate was plotted.

### Flow cytometry

For cell cycle analysis 1 × 10^6^ cells were washed with PBS and fixed with 70% ethanol. Subsequently, they were stained with propidium iodide (BioLegend, Cat # 421301) as per the manufacturer’s protocol before acquisition. Cell cycle distribution was assessed using an inbuilt univariate model of FlowJo software.

For immunophenotyping experiments, 5 × 10^5^ cells were stained using CD117-APC (Miltenyi Biotec, 130-098-207, 1:50) and CD133-PE (Miltenyi Biotec, 130-113-670, 1:50) antibodies for 1hr on ice. Next, data was acquired, and gates were applied to identify positive cell populations. All the flow cytometry experiments were performed on BD FACSMelody™ Cell Sorter or The BD Accuri™ C6 Plus Flow Cytometer and the acquired data was analyzed using FlowJo v.10.6.1.

### Correlation analysis

Compiled colorectal cancer data from different publicly available microarray studies (N=1538) and TCGA-COADREAD was used to check the correlation of *DKC1* with genes involved in the DNA repair pathway. R package ‘corrplot’ was used for the correlation analysis. The function ‘cor’ was used to calculate the Pearson correlation for each gene with every other gene. The function ‘corrplot’ was used to make a correlogram based on the correlation matrix from the ‘cor’ function.

### Immunofluorescence

2 × 10^4^ cells were plated per coverslip per well of 4-well cell culture dishes. Next day, cells were washed with phosphate-buffered saline (PBS) and fixed with 4% paraformaldehyde in the dark. Cells were washed post-fixation, permeabilized using 0.3% Triton X-100 in PBS, and washed with 0.1% Tween-20 containing PBS (PBS-T). Subsequently, 5% normal goat serum (NGS) in PBS-T was used for masking and was removed an hour later followed by washing with PBS-T. Mouse monoclonal antibody against γ-H2AX (Santacruz, sc-517348) was diluted (1:50) in 2% NGS in PBS-T and cells were incubated in it overnight at 4°C. Following day, the primary antibody was removed, and cells were washed with PBS-T to remove the non-specifically bound antibody. Cells were then incubated for 2 hrs at room temperature with Alexa-fluor-555 conjugated secondary antibody against mouse heavy chain (dilution 1:600) in 2% NGS in PBS-T. Next, the secondary antibody was washed with PBS-T, and cells was incubated with Hoechst nuclear staining dye (5μg/mL) for 45 min. Cells were washed again and coverslips were mounted on slides (Vectashield) followed by imaging using Carl Zeiss Axio Z1 observer epifluorescence microscope with an Apotome device attached.

### Tumor Sphere Assay

Cells were plated in an ultra-low attachment cell-culture dish (Corning) at a density of 1.5 × 10^4^ cells/well and were cultured in DMEM-F12 serum-free medium supplemented with growth factors (EGF and FGF), Insulin, and B-27 (all these from Invitrogen). Cells were allowed to grow, and spheres formed were dismantled by pipetting every third day. The experiment was terminated after three weeks, and spheres were counted and imaged using the brightfield microscope (Leica). The diameter (d) of spheres was quantified using ImageJ and the surface area was calculated using the formula *πd*^2^. Post-image acquisition, RNA was isolated from these tumor spheres for the quantitation of stemness markers.

### Chemosensitivity assay

5 × 10^3^ SW620 control (shSCRM) and *DKC1* knockdown (sh*DKC1*-2) cells were plated in 96 well dish. 5-Fluorouracil concentration in a range of 400μM to 0.39 μM was applied for 48hrs and the media with the drug was replenished once every 24hrs. 10 μl of resazurin cell viability reagent was added to each well after treatment and incubated for 3.5 hrs. Following this, emission was measured at 590nm with excitation at 530nm using the Synergy Z4 fluorescence reader (Biotek, Winooski, USA). The data was analyzed using the GraphPad prism 7.0 and IC_50_ values of the drugs in the cell lines were calculated with the Non-linear regression method.

### Foci formation assay

For foci formation assay, cells (2×10^3^) were plated in six-well culture dishes in cell line-specific recommended culture media supplemented with 2% heat-inactivated fetal bovine serum (Invitrogen) and incubated at 37°C, media was changed on alternate days. The assay was terminated after 10 days and cells were dried and stained with crystal violet solution (0.05% w/v).

### Mice xenograft and immunohistochemistry

NOD.CB17-Prkdc^scid/J^ (NOD/SCID) (Jackson Laboratory) male mice were randomized and used for the subcutaneous xenograft assay. The experiment was performed as per the standard procedure and the guidelines of Committee for the Purpose of Control and Supervision of Experiments on Animals (CPCSEA) were strictly followed. Regulatory standards of the Institutional Animal Ethics Committee of the Indian Institute of Technology Kanpur were precisely maintained during the experiment.

First, Ketamine (50mg/kg) and Xylazine (5mg/kg) cocktail was used to anesthetize the male mice (randomized) via intraperitoneal injection. SW620 shSCRM control, sh*DKC1*-2, and sh*DKC1*-3 cells were trypsinized and 1.5 × 10^6^ cells were resuspended in 100μL of 0.9% saline containing 15% Matrigel and were subcutaneously implanted at the dorsal side on both flanks (N=6 for each type). The mice were monitored for tumor growth on alternate days and tumor sizes were measured using digital Vernier Calipers and volume was calculated with the following formula: (π/6) (L × W^2^), (L = length; W = width).

For immunohistochemical staining, tumors were first fixed in 10% buffered formalin, paraffinized and 3μm sections were taken using microtome (Leica). Tumor xenograft slides were incubated at 60 °C for 2 h followed by antigen retrieval in a PT Link instrument (Agilent DAKO, PT200) using EnVision FLEX Target Retrieval Solution, High pH (Agilent DAKO, K800421-2) for DKC1 slides and EnVision FLEX Target Retrieval Solution, Low pH (Agilent DAKO, K800521-2) for Ki67 slides. 1X EnVision FLEX Wash Buffer (Agilent DAKO, K800721-2) was used to wash slides followed by treatment with Peroxidazed 1 (Biocare Medical, PX968M) and Background Punisher (Biocare Medical, BP974L) for 5 min and 10 min respectively with wash after each step. Slides were incubated with antibodies against DKC1 (Abcam, ab156877) and Ki67 (clone MIB-1, DAKO M7240) overnight at 4°C.

Afterward, slides were washed and incubated in Mach2 Doublestain 1 (DKC1 slides) (Biocare Medical, MRCT523L) or Mach2 Doublestain 2 (Ki67 slides) (Biocare Medical, MRCT525L) for 30 min at room temperature, rinsed in 1X EnVision wash buffer 3 times and treated with a Betazoid DAB solution (Biocare Medical, BDB2004H) for 5 min. Next, slides were rinsed in distilled water followed by treatment with EnVision FLEX Hematoxylin (Agilent DAKO, K800821-2). After rinsing in tap water, slides were dried completely and then dehydrated using xylene. EcoMount (Biocare Medical, EM897L) was added to each slide, which was then mounted with coverslips.

DKC1 quantification was performed by calculating integrated density for 15 random histological fields using the IHC Toolbox plugin of ImageJ software. For Ki67 positive nuclei quantification, a ratio of Ki67 positive to total nuclei was calculated for 15 random histological fields using ImageJS Ki-67 online module ^60^.

Patient tissue microarray (TMA) slides (CO1502, Tissuearray.com LLC) were subjected to immunohistochemical staining as per the mentioned protocol. Except for SGPP2 IHC, where the TMA slide was incubated with antibody against SGPP2 (PA5-42767) overnight at 4°C. SGPP2 slide was washed and incubated in Mach2 Doublestain 1 for 30 min at room temperature. For DKC1 staining of TMA, the same protocol was followed as mentioned above.

### ^6061^In silico analysis for CMS classification

DKC1-high and DKC1-low patients were subjected to Gene Set Variation Analysis (GSVA) for up and downregulated CMS signatures derived from the CMS classification ^43^ or a smaller signature proposed for the same as the ‘ColoType’ ^24^. For proteomics data analysis, Clinical Proteomic Tumor Analysis Consortium (CPTAC) proteomics data for colorectal cancer was utilized to assess different parameters.

### Single-cell expression data analysis

Single-cell expression matrices and metadata were downloaded from GEO datasets GSE132465 and GSE144735. The normalized expression data and metadata were combined into an anndata object using Scanpy. Further analysis and visualizations such as dot plots were done using Scanpy functions.

### siRNA transfection

Cells were plated to ∼50% confluence in 6-well plates and 30pmol of siRNA against *CTNNB1* (si*CTNNB1*) or non-targeting siRNA (siNT) (Dharmacon) were transfected using XtremeGENE™ siRNA Transfection Reagent (Roche, Merck) as per manufacturer’s instructions (for SW620 150pmol siRNA was used). Cells were transfected again after 24 hrs, and protein or RNA was isolated post 36hrs of second transfection and were analyzed by qPCR and immunoblotting as mentioned above.

### WNT3A stimulation

WNT3A-conditioned media was collected as per the ATCC recommendations. Briefly, L Wnt- 3A cells or control LM cells were plated at ∼20% confluency and two batches of conditioned media were collected at the fourth and seventh day, mixed, filtered, and stored at 4°C. SW620 cells in a 100mm dish were serum starved for 6hrs followed by supplementation with 50% conditioned medium. The media was replaced every 24hrs and the experiment was terminated after 60hrs since the start of the treatment.

### Enzyme-Linked Immunosorbent Assay for Sphingosine-1 phosphate

Sphingosine-1 phosphate levels were measured using ELISA as per the manufacturer’s instructions (Universal SIP ELISA kit, Maker: ImmunoTag, G bioscience, ITLK09484). Briefly, cells in 100mm dishes were lysed using a buffer containing 1% Triton X-100, 150mM NaCl, 1mM EDTA, 1mM EGTA, 1mM Na_3_VO_4_, 10mM HEPES (pH 7.5) along with protease and phosphatase inhibitors. Lysis was followed by centrifugation @13000rpm @ 4°C for 10’. The supernatant was collected and the protein amount was estimated using the bicinchoninic acid assay, and was incubated in 96-well strips pre-coated with antibody against S1P, followed by incubation with biotinylated antibody. Next, Streptavidin-HRP solution was used followed by TMB substrate. The reaction was stopped using ‘Stop solution’ and absorbance was measured at 450nm. S1P levels normalized to protein expression were used for the final assessment.

### ChIP-sequencing data analysis

Publicly available ChIP-Seq data (GSE31477) for TCF7L2 in HCT116 cells was used to explore the regulation of DKC1 under the control of WNT signaling. Quality assessment for raw single-end reads was done using FASTQC, trimmed with FASTQ Trimmer, with default settings of Galaxy web platform (usegalaxy.org). Reads were aligned to the reference genome (hg19) using Bowtie to generate Sequence Alignment/Map (SAM) files. Unaligned or unmapped reads were filtered using FilterSAM, a utility of SAMtools. The SAM files were converted to its Binary Alignment/Maps (BAM) files using SAMtools. Further, ChIP-Seq peaks for RNA PolII and TCF7L2 were called using Model-based analysis of ChIP-seq (MACS; P < 10−5) with default settings against Input. BAM and BED files obtained were visualized by Integrative Genomic Browser (IGB).

### Immunoprecipitation

SW620 cells were cultured in a 100mm dish and were harvested at 90% confluence; cells were washed twice with PBS and lysed with ice-cold lysis buffer containing NP-40 (2%, ab142227, Abcam), Triton X-100 (2%), NaCl (100mM), Tris-Cl (50 mM, pH 8.0), EDTA (1 mM), 1X protease inhibitor (Genetix) and 1X phosphatase inhibitor (524625, Millipore) and kept at 4°C on rotor for 1hr, centrifuged @ 13000rpm @ 4°C for 10mins. The supernatant was incubated with 4µg of primary DKC1 (Santacruz, sc-373956-X, mouse monoclonal antibody) or isotype control antibody (mouse IgG, Invitrogen, 10400C) at 4°C overnight. Simultaneously, protein A/G coated agarose beads (SantaCruz, sc-2003) were blocked with bovine serum albumin (HiMedia) followed by incubation at 4 °C overnight. Blocked beads were incubated for 24hrs at 4°C with the lysate to make antibody-bead conjugates. Next, these conjugates were washed and immunocomplex was eluted using elution buffer (2X dye containing SDS).

### Chromatin Immunoprecipitation

ChIP experiments were performed as previously described ^61^. Briefly, SW620 cells or SW620 scrambled control and *DKC1* knockdown cells were plated in a 100mm cell culture dish and were harvested at 90% confluence. First, cells were fixed with 1% formaldehyde for 10 mins, quenched with 125mM glycine, washed with PBS, and lysed using lysis buffer containing 1% SDS, Tris-Cl (50 mM, pH 8.0), EDTA (10 mM) and 1X protease inhibitor (Genetix) followed by DNA fragmentation through sonication to an average fragment length of ∼500 bp using Bioruptor (Diagenode) and sheared chromatin was incubated at 4°C overnight with 4µg of primary β-catenin (CST, D10A8) or SOX2 (Abcam, ab97959), or isotype control antibodies (rabbit IgG, Invitrogen, 10500C). Concomitantly, Protein G-coated Dynabeads (Invitrogen) were blocked with bovine serum albumin (HiMedia) and sheared salmon sperm DNA (Sigma-Aldrich), followed by incubation at 4 °C overnight. Blocked beads were incubated for 8hrs at 4°C with the lysate to make antibody-bead conjugates. Next, these conjugates were washed and immunocomplex was eluted using elution buffer [1% SDS, 100 mM NaHCO_3_, Proteinase K (Sigma-Aldrich) and RNase A (500 µg/ml each) (Sigma-Aldrich)]. Phenol-chloroform-isoamyl alcohol extraction method was used to isolate DNA from the immunocomplex. JASPAR and MatInspector softwares were used to predict binding sites for β-catenin/TCF complex on the DKC1 promoter or SOX2 on the SGPP2 promoter.

### Lipidomics analysis

To analyze the global lipid profiles, patient tumor specimens were homogenized in 150 mM Ammonium bicarbonate solution in a 1:10 ratio (w/v, mg/uL) with a Tissuelyser system, 2 cycles for 2 min at 28hz frequency @4 degrees with one 5mm stainless steel bead followed by sonication for lysis. Following this, 200uL of each tissue homogenate was taken from every sample, and UHPLC grade methanol was added to it in a 1:5 ratio (v/v, uL/uL) and snap frozen using liquid nitrogen stored at -80°C. Similarly, for SW620 control and *DKC1* knockdown cells, ∼15 million cells were used in triplicates. The cells were detached using trypsin, neutralized and supernatant was removed and resuspended in 150mM Ammonium bicarbonate. Next, lysis was performed using sonication and protein concentration was estimated directly for each homogenate using the BCA method. Cells were again centrifuged and supernatant was removed fixed with 100μl of 100% MeOH, snap frozen in liquid nitrogen and immediately stored at -80°C. The samples were shipped to the Singapore Lipidomics Incubator, National University of Singapore where they were processed further for downstream analysis.

Briefly, 100μl chilled butanol containing internal standards was added to the samples, vortexed for 2 min, shaken, and sonicated at 4°C for 30 min, respectively. Centrifuged twice at 14000 rpm, 4°C for 10min, and the supernatant (BuMe) was collected for LC-MS analysis. For UHPLC: Mobile phase A: 40% acetonitrile + 60% water + 10 mM ammonium formate and Mobile phase B: 10% acetonitrile + 90% isopropanol + 10 mM ammonium formate with Column: Poroshell 120 C18 (2.1 x 50 mm, 1.9 µm) were used at a column temperature of 40°C.

LC Gradient

| Time [min] | Eluent B% | Flow rate [mL/min] |
| --- | --- | --- |
| 0 | 20 | 0.4 |
| 2 | 60 | 0.4 |
| 12 | 100 | 0.4 |
| 14 | 100 | 0.4 |
| 14.01 | 20 | 0.4 |
| 15.8 | 20 | 0.4 |

The positive ionization spray voltage and nozzle voltage were set at 3,000 V and 1,000 V, respectively. The drying gas and sheath gas temperatures were both maintained at 250 C. The drying gas and sheath gas flow rates were 14 L/min and 11 L/min, respectively. The nebulizer nitrogen gas flow rate was set at 35 psi. The iFunnel high and low-pressure RF were 150 V and 60 V, respectively. Quantification data was extracted using Agilent MassHunter Quantitative Analysis (QQQ) software. The data was manually checked to ensure that the software integrated the right peaks.

QC filtering is based on raw peak areas for all the measured lipid species using the (pooled) TQC samples. The filtering criteria were as follows:

> Low Signal variation (CoV < 20%);
>
> High Sample/Blank ratio (> 10);
>
> Good Signal linearity (R^2^ > 0.8).

The lipid species which didn’t pass QC filtering are marked as “FAILED” and excluded from the downstream analysis.

Using lipid class specific standards, the relative concentration of an analyte in the sample calculated as:

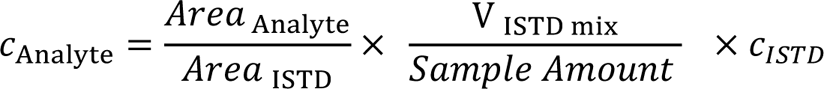

***c_Analyte_* :** Molar [M or µmol/µL] concentration of the analyte in the sample,

*Area_Analyte_* : Peak area of Analyte,

*Area_ISTD_* : Peak area of ISTD,

*V_ISTD mix_* : Extraction solvent volume (ISTD mixture)

*Sample_Amount* : Quantity of the sample (*e.g.* µL or mg)

*c_ISTD_* : Concentration of the ISTD in the extraction solvent (ISTD mixture)

### Multivariate analysis

Principal component analysis (PCA) and Orthogonal Projection of Latent Structures- Discriminant Analysis (OPLSDA) providing Variable Importance in Projection (VIP) scores for dysregulated lipid species were performed using ‘Statistical analysis (one factor)’ module of Metaboanalyst v5.0.

### Integrative pathway analysis

Differential genes and lipids obtained from RNAseq and UHPLC-MS/MS analysis for scrambled control and *DKC1* silenced SW620 cells were subjected to the Joint-pathway analysis module of Metaboanalyst v5.0. Enrichment analysis was performed using a hypergeometric test and degree centrality was used as the topology measure to combine queries. The analysis was performed to identify significantly impacted metabolic pathways.

### Biomarker Analysis

The lipid profile of CRC patient tumors was used to identify biomarkers using the Biomarker Analysis module of Metaboanalyst v5.0. The data was Pareto scaled for normalization. Classical univariate analysis was performed to generate Receiver Operating Characteristic (ROC) curve and compute the area under the curve for different ceramides. ROC curve-based model evaluation (Tester) module was used to assess the effectiveness of proposed ceramides as biomarkers for *DKC1*-High tumors.

### Statistical Analysis

Each experiment was performed atleast in triplicates using multiple biological replicates. GraphPad Prism 7.0 was utilized for statistical analysis. For all comparisons, two tailed Student t-tests or One-Way ANOVA were used unless otherwise stated in respective figure with P-values less than 0.05 considered as significant.

## Acknowledgments

We are grateful to Dr. Eric R. Fearon, University of Michigan, Ann Arbor, for providing WiDr colorectal cancer cell line. We also thank Dr. Jonaki Sen and Dr. Pradip Sinha for extending the use of the microscopy facility. The Proteome data used in this publication were generated by the National Cancer Institute Clinical Proteomic Tumor Analysis Consortium (CPTAC). We also thank the Singapore Lipidomics Incubator, the National University of Singapore for extending the use of their UHPLC-MS/MS facility for lipidomics.

## Funding

Science and Engineering Research Board (SERB)-POWER grant (SPG/2021/000851) to BA. India Alliance DBT/ Wellcome Trust Senior Fellow (IA/S/19/2/504659) to BA.

Science and Engineering Research Board, Ministry of Science and Technology, Government of India (EMR/2016/005273) to BA.

S. Ramachandran-National Bioscience Award for Career Development (BT/HRD/NBA/NWB/39/2020-21) by the Department of Biotechnology (DBT) to BA.

## Author Contributions

SN and BA conceptualized the study. SN, AP, SC, NP are responsible for the methodology. Investigation & Analysis were performed by SN, AP, UK, AS, SC, NP. Patient samples were collected by SN with the help of AK, AT, AC. Supervision, funding acquisition, project administration were done by BA. Original draft was written by SN, BA. Review & editing were done by SN, AP, UK, BA.

## Data and materials availability

RNAseq expression data will be deposited on the GEO database. Lipidomics data will be submitted to UCSD Metabolomics Workbench. R codes are publicly available for tools used here for *in silico* analysis.

