## Supplementary Figure 1 to 5 for "Transcriptional regulation of Dyskerin *via* canonical WNT signaling modulates sphingolipid biosynthesis and drives colorectal cancer"

**Affiliations:**

### Supplementary figure 1

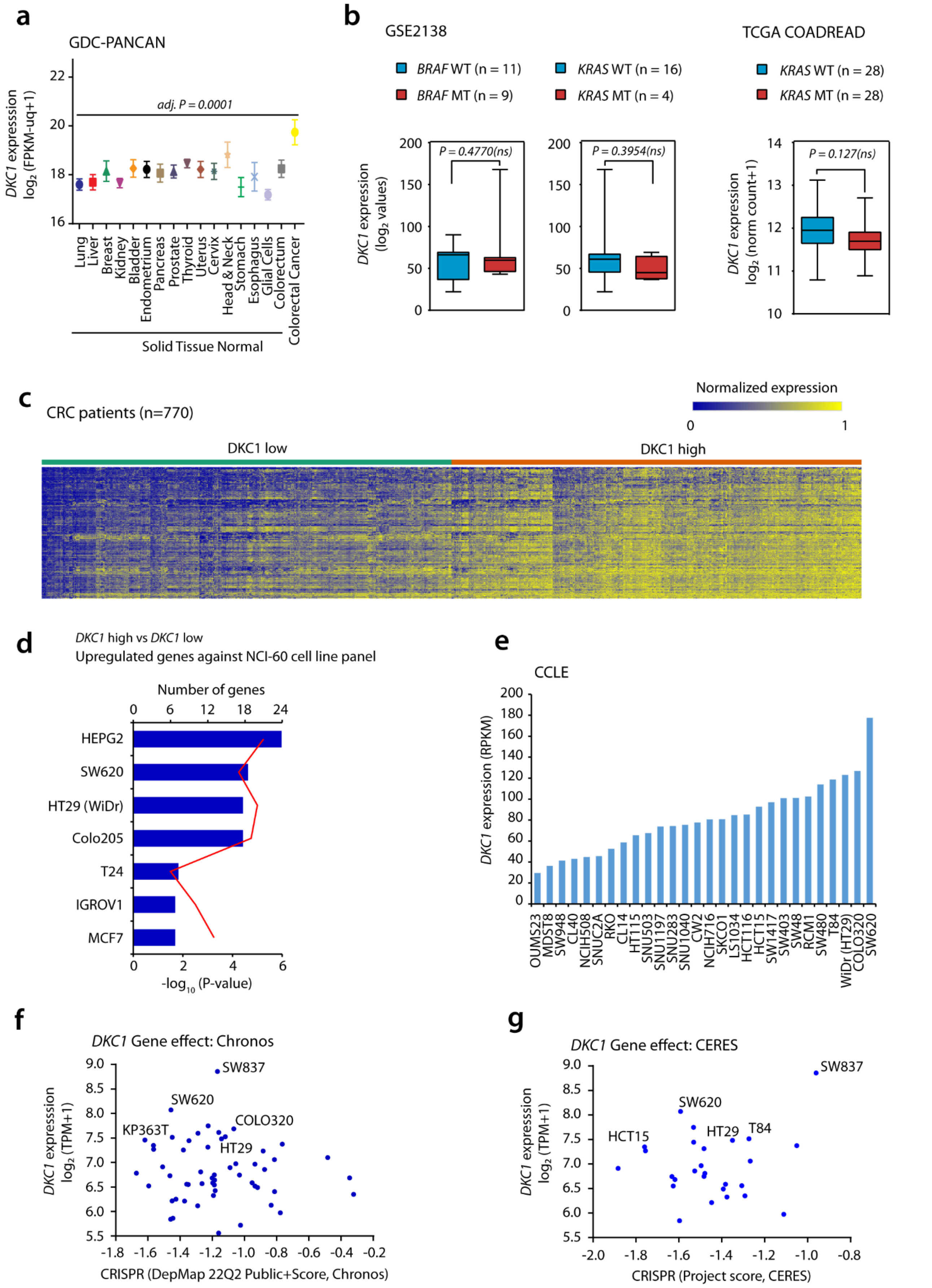


**Fig. S1. High DKC1 expression associates with a significant fraction colorectal cancer patients. a,** DKC1 expression in normal tissues and CRC tumors from Genomics Data Commons Pan-Cancer (GDC-PANCAN) atlas. **b**, DKC1 expression in KRAS/BRAF mutant (MT) and wild type (WT) CRC cases. **c,** Heatmap depicting association of upregulated genes obtained using gene expression data from DKC1-high vs DKC1-low cases (N=770). **d,** Enrichment analysis of the same genes as in **c** against the NCI-60 cell line panel using EnrichR. **e,** Expression of DKC1 in different colorectal cancer cell lines. **f and g,** Dependency of CRC cell lines in publicly available Chronos and CERES studies from DepMap portal of CCLE. For **a and b**, Significance was evaluated using two tailed Student's t-test. Data are expressed as mean ± s.e.m.

### Supplementary figure 2

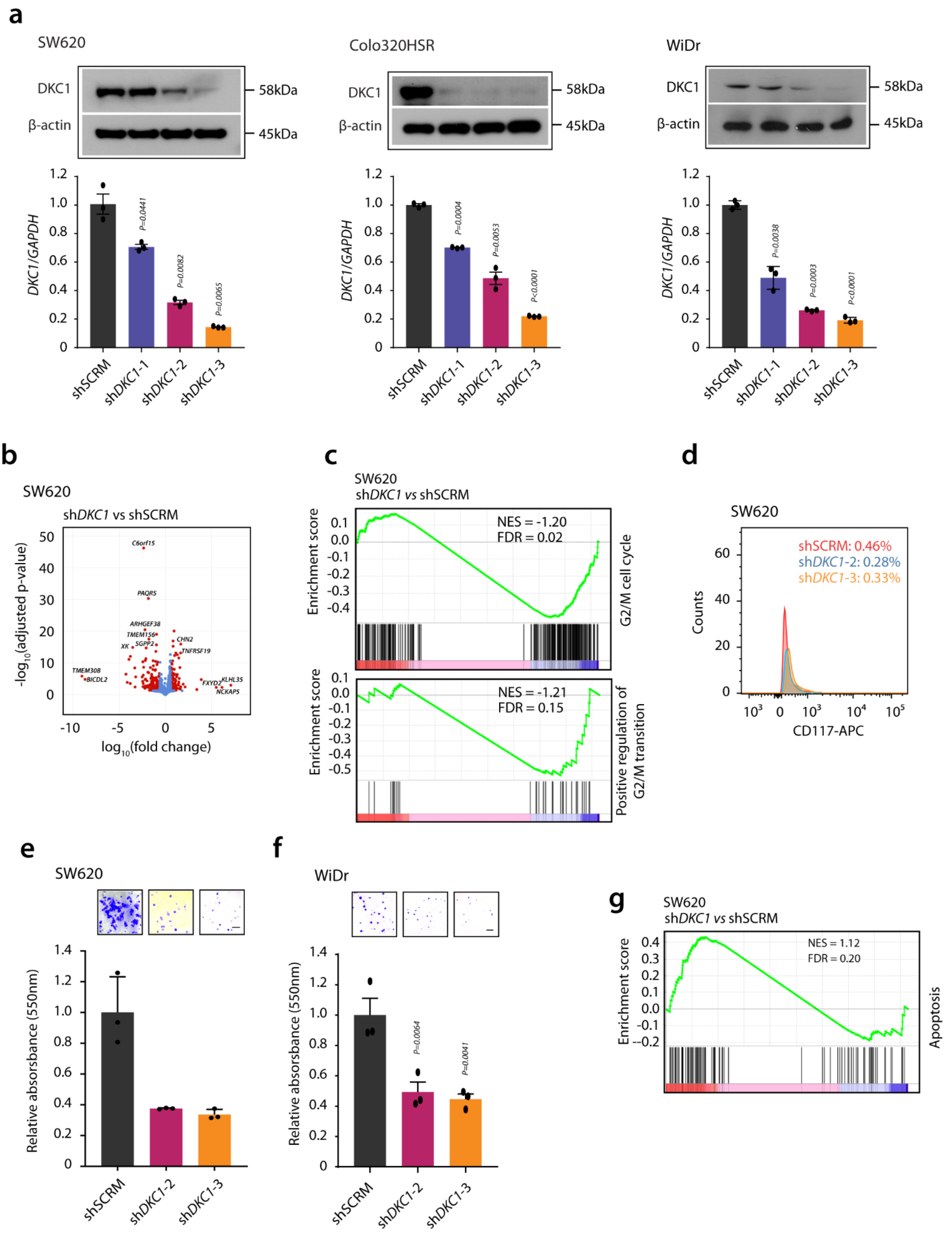


**Fig. S2. High DKC1 promotes oncogenicity in CRC. a,** Transcript and protein levels of DKC1 in SW620, Colo320HSR and WiDr cells. **b,** Volcano plot showing differentially expressed genes from RNAseq analysis of SW620 DKC1 knockdown and control cells. **c,** Gene Set Enrichment Analysis (GSEA) of the gene expression data from same cells as in **b** for G2/M cell cycle transition pathways. **d,** Flow cytometry-based cell surface expression of CD117 labeled with APC in same cells as in **b**, graphs represent percent positive cells. **e and f,** Foci formation assay for SW620 and WiDr DKC1 knockdown and control cells (scale bar = 1mm). **g,** GSEA of the gene expression data for apoptosis pathway from same cells as in b. For **e and f,** Unpaired Student's two-tailed t-test with Welch's correction were applied. Data are expressed as mean ± s.e.m.

### Supplementary figure 3

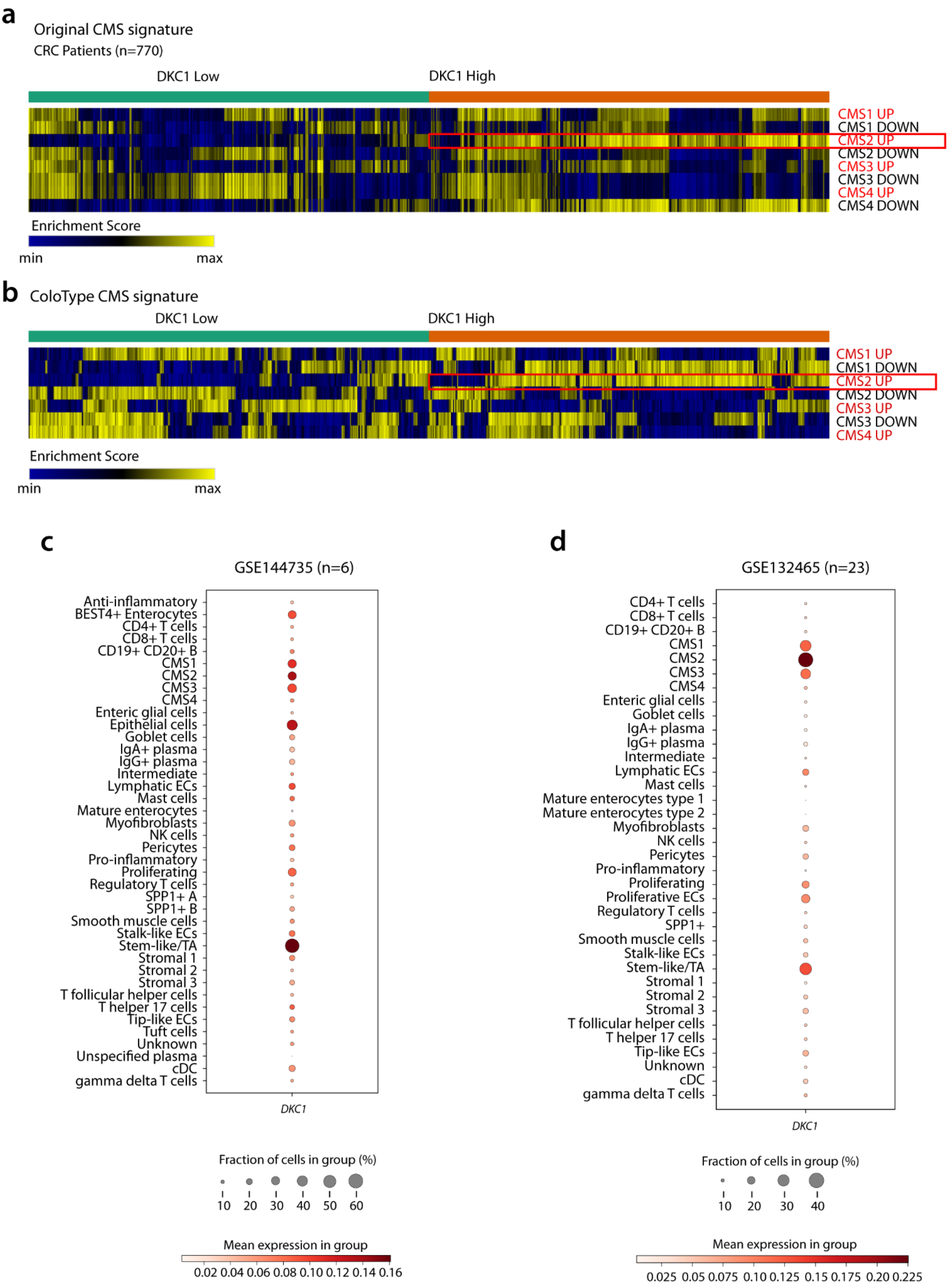


**Fig. S3. High DKC1 expression associates with CMS2 subtype of CRC. a,** Heatmap depiction of Consensus Molecular Subtype (CMS) gene signatures with gene expression data from DKC1-high vs DKC1-low cases (N=770) using Gene Set Variation Analysis (GSVA), DKC1-high patients showing association with 'CMS2 UP' gene signature is marked in red box. **b,** Same as in a except a smaller 40 gene signature 'ColoType' for CMS classification was used here for GSVA. **c and d**, Heatmap depiction of single cell expression data from patients of Belgian (GSE144735, N=6) and Korean (GSE132465, N=23) ethnicities respectively, showing the fraction of cells and mean expression of DKC1 in each group.

#### Supplementary Figure 4

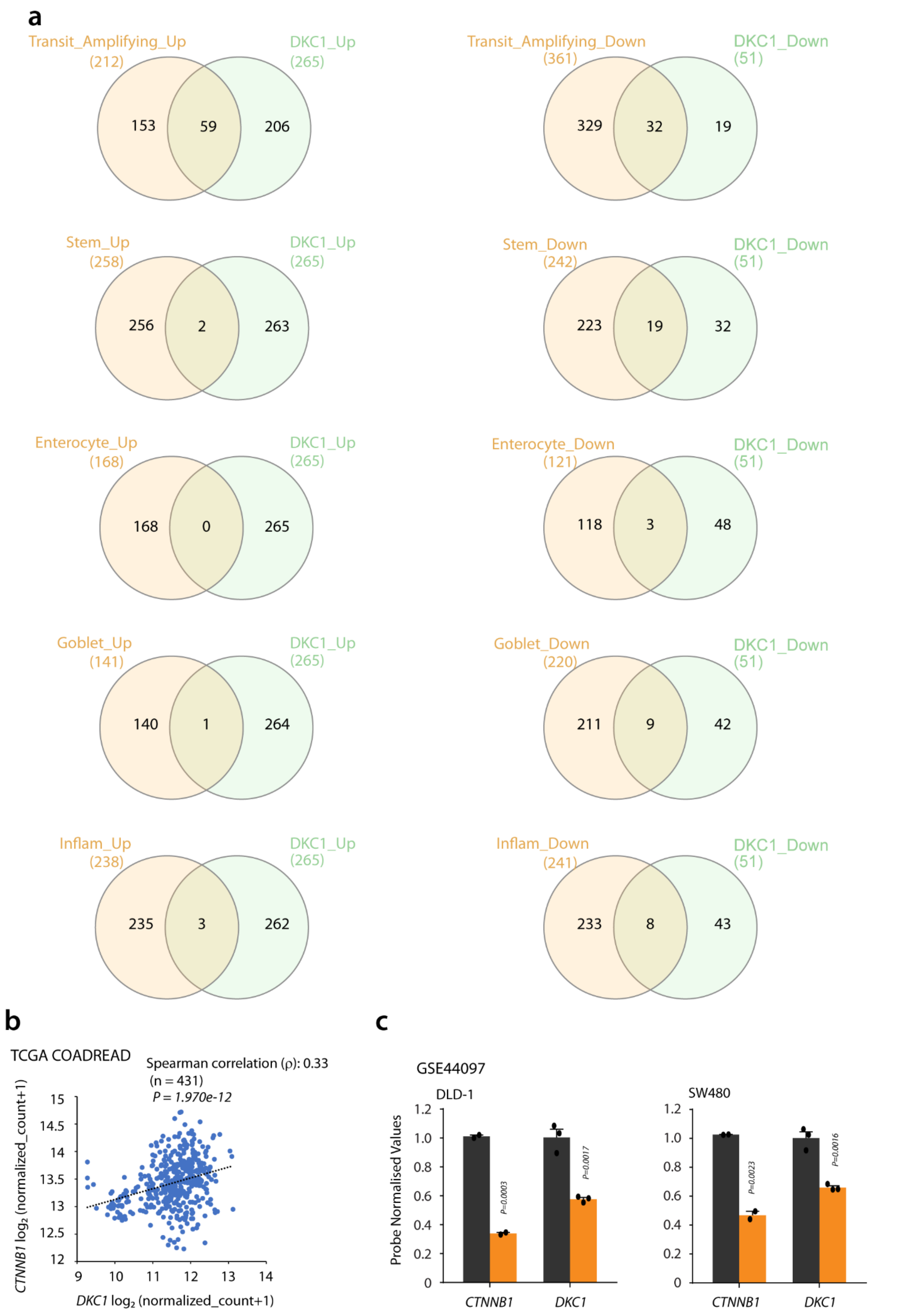


**Fig. S4. DKC1 associates with Transit Amplifying/stemlike subtype and WNT signaling in CRC. a,** Venn diagrams showing overlaps between cell-type based gene signatures and upregulated genes obtained using gene expression data from DKC1-high vs DKC1-low cases (N=770). **b,** Scatter-plot depiction of correlation between DKC1 and CTNNB1 in gene expression data of primary tumor and adjacent normal tissues from TCGA COADREAD dataset. **c,** DKC1 expression from microarray analysis of CTNNB1 silenced and control DLD1 and SW480 cells ( GSE44097). For panel **c,** Significance was evaluated using two tailed Student's t-test. Data are expressed as mean ± s.e.m.

### Supplementary figure 5

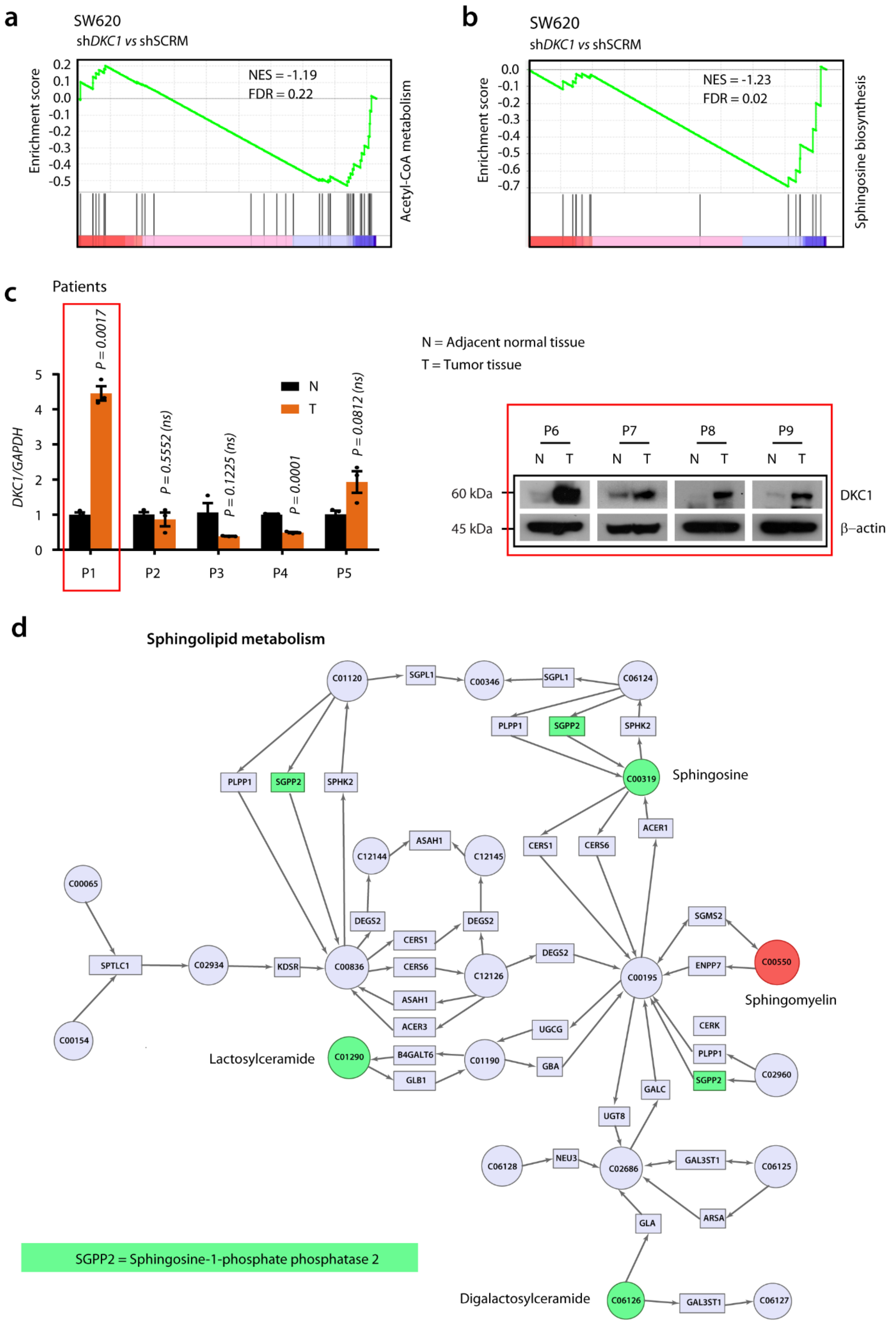


**Fig. S5. Sphingolipid biosynthesis is dysregulated in CRC with high DKC1 expression. a and b,** Gene Set Enrichment Analysis (GSEA) of the gene expression data from SW620 DKC1 knockdown and control cells for Acetyl Co-A metabolism and sphingosine biosynthesis respectively. **c,** Transcript (P1, P2, P3, P4, P5) and protein (P6, P7, P8, P9) levels of DKC1 in tumor tissues with respect to adjacent normal tissues, tumor specimens classified as DKC1-high are marked in red boxes. **d,** Detailed map of sphingolipid metabolism obtained from integrative joint pathway analysis using RNAseq and lipid profile data from same cells as in a, rectangles represents genes (official gene symbols) and circles represent metabolites (KEGG codes), red and green colors depict upregulated and downregulated hits. For panel b, significance was evaluated using two tailed Student's t-test. Data are expressed as mean ± s.e.m.
